# Learning new perceptual skills: Individual differences in the computations that integrate novel sensory cues into depth perception

**DOI:** 10.1101/2025.03.25.645045

**Authors:** Meike Scheller, Stacey Aston, Heather Slater, Marko Nardini

## Abstract

Sensory substitution and augmentation rely on the brain’s ability to integrate novel sensory cues into its perceptual repertoire. However, the flexibility of the computations that support augmented perception is still not fully known. Here, we contrasted how a novel depth cue is processed, as compared with familiar depth cues, following one hour of training. Observers (N=78) made forced-choice comparisons of surface distances (depths), while we assessed three markers of integration with familiar and newly learned cues: (1) cue combination, predicting precision benefits; (2) re-weighting, predicting reliability-weighted biases; and (3) congruence sensitivity, predicting increased sensitivity for the learned cue mapping. We found that, (1) while familiar cues (size and binocular disparity) were combined near-optimally, there was little evidence for a novel cue (auditory pitch) being combined with binocular disparity. Measures of repeatability across two sessions suggest that this was due to reliable inter-individual differences in combination with the novel cue. In contrast, (2) both familiar and novel cue pairs were re-weighted by their relative reliabilities and (3) showed sensitivity to incongruence. These direct comparisons show that, while a novel cue was rapidly mapped onto depth and weighted in line with its relative reliability, it was not integrated into the native perceptual repertoire by everyone. Reliable individual differences suggest that abilities to combine novel sensory cues may vary between people. These findings provide insight into the flexibility of human sensory processing and suggest that learning to interpret new sensory information depends in part on individual flexibility in perceptual computations.

**Highlights:**

- We evaluated evidence for integration of novel and familiar cues to depth
- After short training, direct comparisons show dissociations in integration
- At a group level, novel cues, unlike familiar, were not combined overall
- Strong individual differences suggest varied abilities to combine new cues
- Individual differences seem to play an important role in integration of new cues

## 1. Introduction

The brain’s ability to learn which sensory information should be combined into a unified percept fundamentally underpins human perception and cognition. It plays a pivotal role in shaping how we understand our environment and make efficient decisions based on the information we have at our disposal. In the present study, we investigate the ability to efficiently adopt new sensory information into the perceptual repertoire, using well-established paradigms in perceptual integration research.

When multiple independent sensory estimates (cues) convey the same feature information (e.g., visual and auditory cues to location), the adult brain combines them by their respective reliabilities, leading to an increase in perceptual precision. This process has frequently been described as “optimal”, as it often allows observers to attain the maximal possible precision benefit from the available information (Alais & Burr, 2004; Ernst & Banks, 2002; Fetsch et al., 2009; Hillis et al., 2004). Experimentally, this precision increase is considered a key marker showing that cues have been averaged (combined; *cue combination*), measured by decreases in sensory noise (increases in precision) when two cues are available compared to when only the best single cue is available (Ernst & Banks, 2002; Rohde et al., 2016; Scheller & Nardini, 2023). For instance, in their seminal work, Ernst and Banks (2002) showed that, when estimating an object’s size, adults average visual and haptic cues while weighting them by their relative reliabilities, leading to a precision enhancement compared with the individual cues. These precision enhancements were well-predicted by a model of statistically optimal estimation under uncertainty, that weights the available estimates by their relative reliabilities. In a similar manner, many other pairs of redundant cues are combined to generate a more precise representation of the feature of interest - not only between sensory modalities (e.g. vision and audition combined to judge location; Alais & Burr, 2004; Bruns, 2019), but also within (e.g. visual binocular disparity and visual motion parallax combined to judge depth (Bradshaw & Rogers, 1996; Hillis et al., 2004; Landy et al., 1995).

Because the environment we live in is constantly changing, perception needs to retain a certain degree of flexibility. In line with this, human cue combination is intrinsically flexible: it allows the brain to use sensory information efficiently across different contexts to optimize perceptual precision. One way in which this is illustrated is in the dynamic re-weighting of cues as their reliabilities change (Alais & Burr, 2004; Ernst & Banks, 2002). For instance, when the relative sensory reliability of a visual and an auditory cue to location is impacted by visibility conditions, such as in foggy or dimly lit environments, vision may convey less reliable information than audition. As a result, the optimal observer (and humans, in line with this) reduce their relative weighting for the visual cue. This dynamic “re-weighting” of sensory information has been evidenced across a range of sensory modalities and contexts (e.g., Alais & Burr, 2004; Ernst & Banks, 2002; Nardini et al., 2010). Another way in which cue combination offers flexibility in perception is evidenced in the ability to learn new cue mappings. That is, based on statistical co-occurrences in the environment, the brain likely learns which cues go together and provide redundant information about the same environmental feature (Ernst, 2007; Parise et al., 2013, 2014; Parise & Ernst, 2024). This is crucial in order to use information redundancy between cues and distil it into a more precise representation about the common environmental feature. Whether cues are treated as redundant (arising from the same environmental feature) and combined depends on the cues’ spatio-temporal co-occurrence (Körding et al., 2007; Parise et al., 2013, 2014; Shams & Beierholm, 2022) as well as individual experience with sensory statistics (e.g. Bruns & Röder, 2023; Ernst, 2007). That is, through extensive experience in childhood, our brains learn which and how cues map onto the same environmental feature (e.g., Gori et al., 2010, 2012; Kolarik et al., 2013; Scheller et al., 2020), for instance visual size and haptic weight (Charpentier, 1891; Murray et al., 1999) or auditory pitch and visual elevation (Parise et al., 2014). When two sensory cues are assumed to arise from the same environmental feature, they are typically combined into a single, perceptual estimate (Ernst, 2007; Körding et al., 2007).

As described above, *combination* and *dynamic reweighting* of redundant noisy cues have been consistently observed in adults using familiar sensory cues. Developmental studies, however, indicate that these abilities are not innate and typically do not fully mature until middle or late childhood, often after the age of 8-13 years, depending on the senses involved and task at hand (Gori et al., 2008; Nardini et al., 2008; see Figure 7 in Scheller et al., 2020, for a summary). As combination is perceptually beneficial, this raises an important question: do these processes depend on the maturation of cognitive functions or do they merely require sufficient experience with correlated sensory cues? More importantly, is the flexibility of the brain to learn and process novel sensory cues in the same way that other, familiar cues are processed constrained to development, or does it extend into adulthood? This question is not only of theoretical importance for understanding the flexible and adaptive nature of the learning and developing brain, but also allows us to better understand the extent to which humans can adopt novel sensory capabilities, such as perceiving the environment though augmented senses or substituting damaged or impaired senses in adulthood (Bach-y-Rita & W. Kercel, 2003; Kärcher et al., 2012; Nardini et al., 2024).

Using cue weighting and perceptual precision improvements (indicative of cue combination) as markers of native perceptual functioning, recent research demonstrated that even with very little training, adults can quickly learn to deploy novel cues to efficiently localize positions in space, a fundamental skill for interacting with our environment (Aston, Beierholm, et al., 2022; Gibo et al., 2017; Negen et al., 2018). For instance, participants were trained to associate auditory click-delay cues with the visual cues to distance of a virtual avatar. After only 600 trials of associative learning (with feedback), distance perception with novel and familiar cues was more precise with both cues together, compared to the best individual cue, and the cues were dynamically re-weighted in accordance with the relative single cue noise (Negen et al., 2018, 2023). Similarly, novel mappings of arbitrary visual cues (colour, shape, height and angle) were combined with visual spread cues to enhance precision in visual, horizontal localization after only 72-216 trials of training (Aston, Beierholm, et al., 2022). These immediate effects suggest that the brain retains a high degree of flexibility to learn to integrate novel cues into native perception. However, novel cues were still treated differently compared to how familiar cues would be treated according to the statistical optimality model. That is, across all three studies, the precision increase, evidencing cue combination, was not as efficient as would have been predicted by reliability-based weighting (Aston, Beierholm, et al., 2022; Negen et al., 2018, 2023). Instead, it was consistently sub-optimal. Furthermore, novel cues were not automatically fused with existing, familiar cues, indicated by an insensitivity to mapping incongruence (Aston, Pattie, et al., 2022; Negen et al., 2023). In contrast, a previous study that asked adults to learn an arbitrary mapping of visual brightness to haptic stiffness cues found enhanced sensitivity to incongruence after only 500 trials (Ernst, 2007). Hence, the extent to which mapping conflict (congruence) affects how cues are dealt with when they are familiar vs novel.

To better understand the scope of the adult brain to adopt new sensory skills, we need to understand the computations that lie at the foundation of sensory combination and provide the plasticity and flexibility to use new sensory cues. Results so far suggest that after short learning, cue combination computations with new cues are fundamentally different to those with highly familiar cues. There has been evidence for precision gains and reweighting consistent with combination, yet the combination process has been consistently less efficient than optimal, unlike in classic cue combination experiments with familiar cues (Aston, Beierholm, et al., 2022; Aston, Pattie, et al., 2022; Negen et al., 2018, 2023; but see also Rahnev & Denison, 2018). However, the way in which combination with novel vs familiar cues really differs is currently unclear. No study to date has directly shown this difference between familiar and novel cue combination, because the conclusions have come from different participants, stimuli, and experiments. To that end, the present study directly compares how familiar and novel cues are combined to enhance perceptual precision, re-weighted, and affected by mapping conflict, using the same distance discrimination tasks and participants. By providing only little training experience with the novel cue (336 trials), we can better understand the immediate flexibility of the different computations underlying cue combination that allow the instant deployment of novel sensory abilities. To better understand the extent to which the adult human brain is able to integrate completely novel cues for the perception of spatial location (depth), the present study tested the flexibility of integration of depth cues at different stages:

○ *Cue combination*, indicated by a precision enhancement when two cues are available compared to the best single cue. This is the classic hallmark of integration into a coherent representation.
○ *Dynamic sensory re-weighting* in accordance with the reliability of the sensory information, indicated by a reduction in relative weight for the cue in which sensory noise was added.
○ *Sensitivity to mapping incongruence*, indicated by the reduction in sensory precision when cues are presented simultaneously in an incongruent (opposite to learned) vs congruent (learned) fashion.

By directly comparing how familiar-familiar cue pairs and familiar-novel cue pairs are processed we will gain a better understanding of the flexibility with which the adult human brain incorporates new sensory experiences and adopts novel sensory cues into its perceptual repertoire. Furthermore, the present study design generates a testbed to ask further questions about the neural processing basis and subjective experiences underlying human cue learning and integration.

## 2. Methods

### 2.1. Experiment overview

This study addresses seven primary research questions, from which testable hypotheses were derived (Table 1). These were categorized into three different markers of native multisensory perception: *combination, re-weighting*, and *incongruence sensitivity* (see Figure 1), one for each type of cue pairings (familiar-familiar; familiar-novel), as well as a final question that contrasted all three markers between the cue pairings.

**Table 1:**
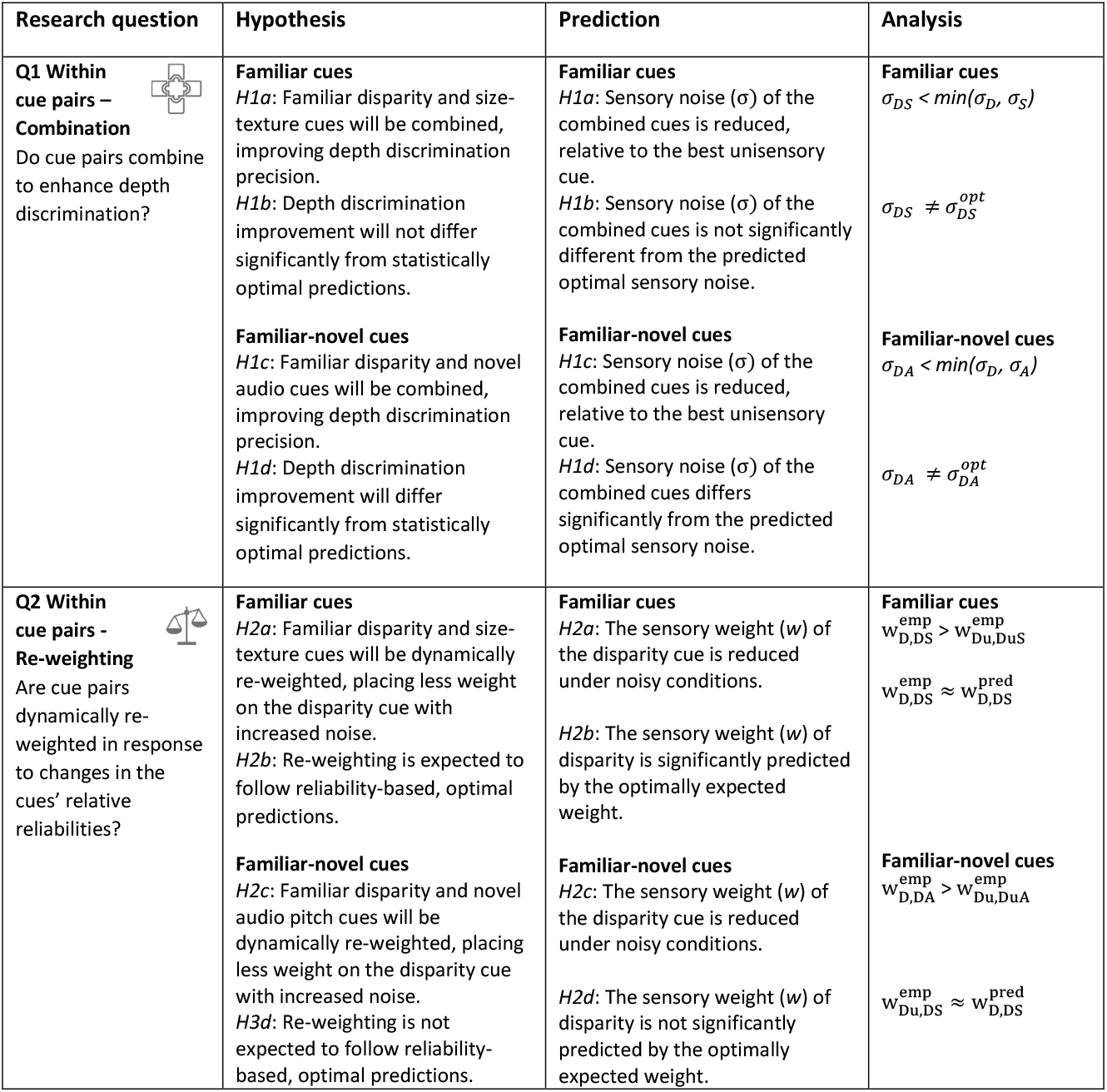
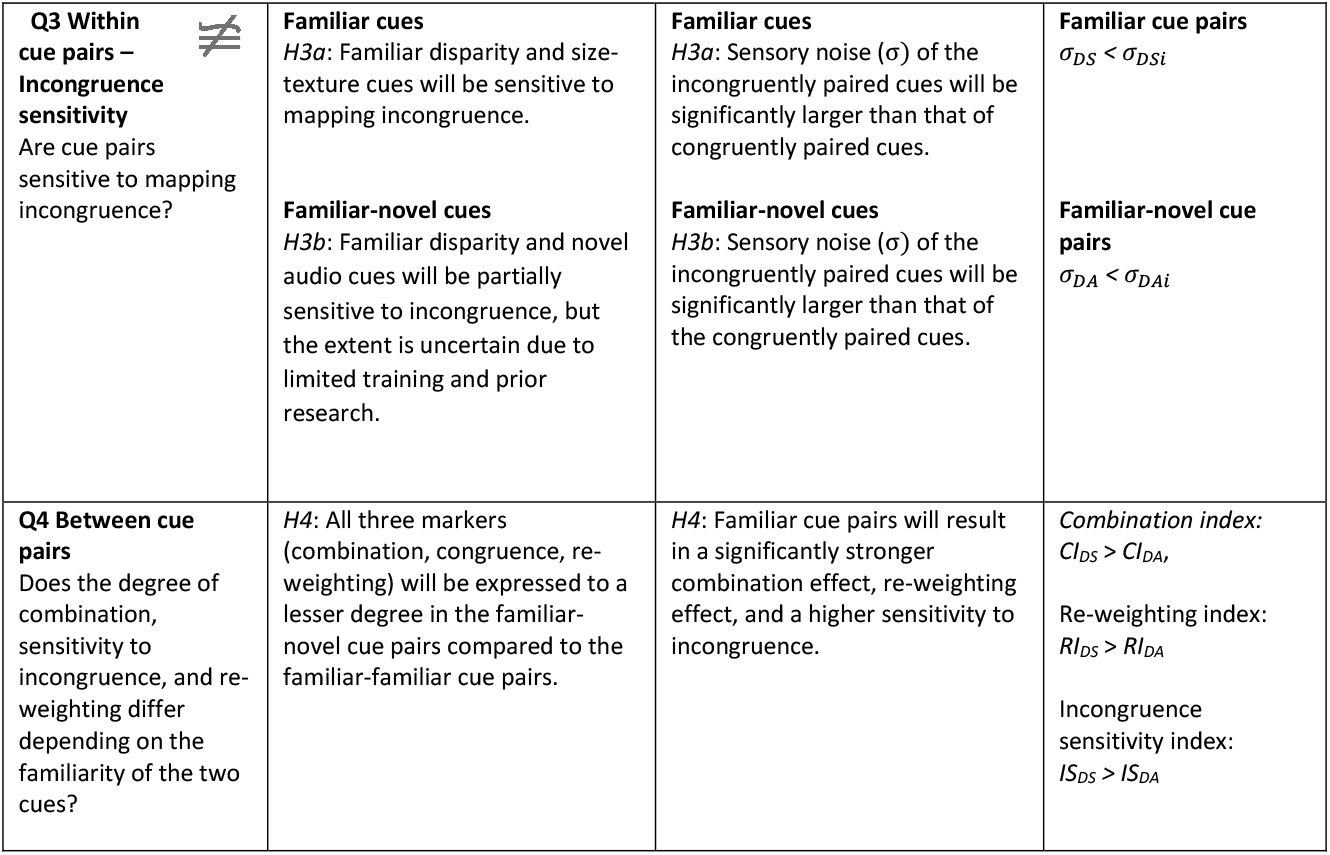
Research questions, hypotheses, and predictions, alongside associated analyses. Research questions one to three, each indicating one marker of perceptual functioning, were addressed for each cue pair (familiar-familiar, familiar-novel), separately. Research question four directly contrasts the two cue pairs on each marker. σ = sensory noise; D = disparity cue alone; S = size cue alone, A: audio cue alone; DA: disparity and audio cues together; DS: disparity and size cues together; opt = optimal prediction; 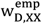 = relative sensory weight for disparity cue, empirically measured, in condition XX; 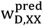 = relative sensory weight for disparity cue, predicted from individual cues, in condition XX

**Figure 1:**
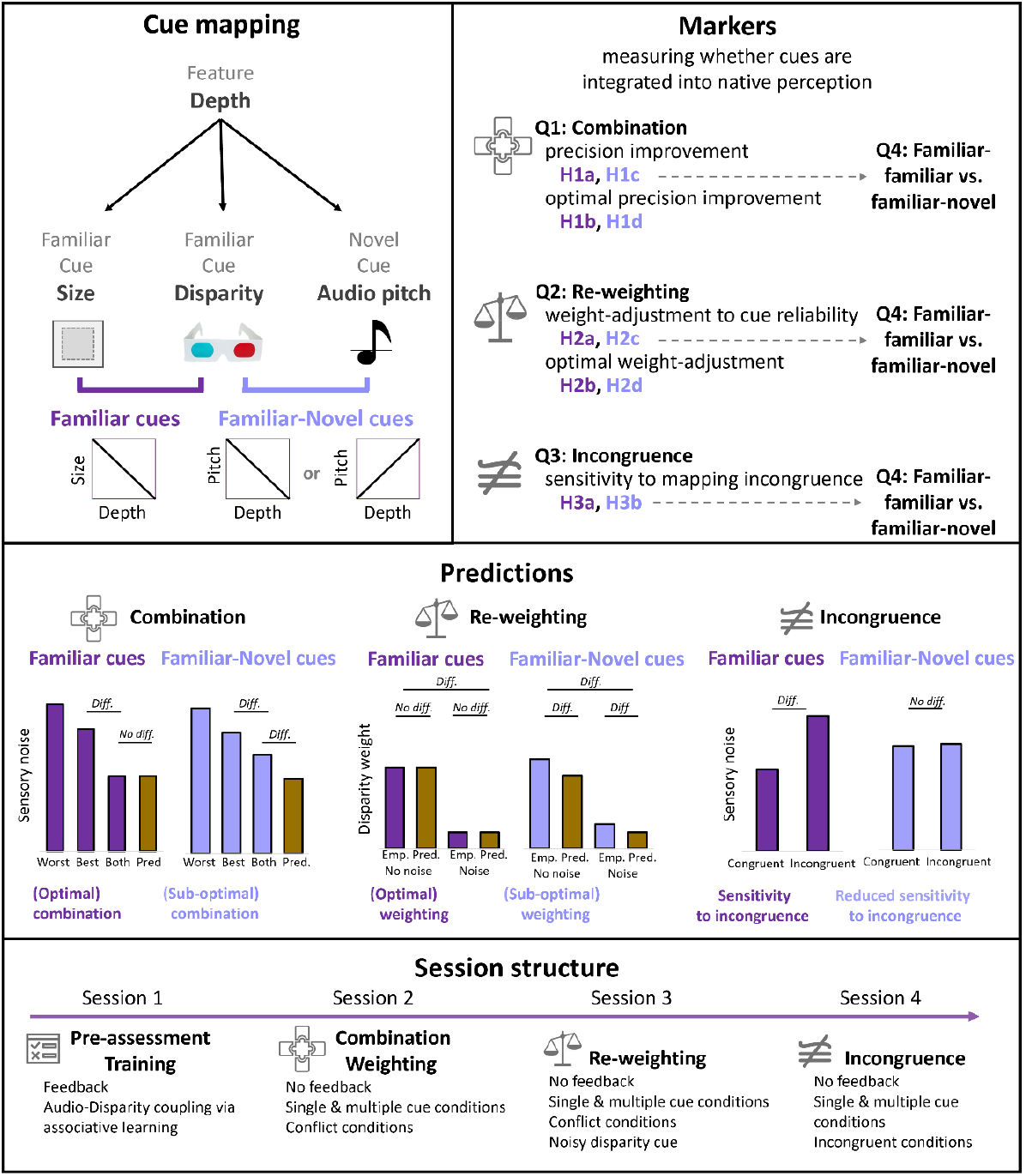
***Top* l*eft***: Visual schematic of familiar and novel cue mappings. Changes in depth were conveyed via binocular disparity cues, size cues, which were following a physically veridical inverse relationship, and novel audio cues. The mapping of the audio cue to depth was counterbalanced across participants, with some learning a positive mapping (pitch increases with depth) and some learning a negative mapping (pitch decreases with depth). ***Top right***: Outline of the three markers that were tested to assess the extent to which novel cues are integrated into native perception. Each marker is tied to a different research question and assessed for both familiar-familiar (dark purple) and familiar-novel (light purple) cue pairs. The contrast between the cue pairs is assessed in Q4 for each marker. ***Middle***: Predictions for the different markers and cue pairs. Combination is indicated by a decrease in sensory noise for the cue pairs compared to the best single cue. Optimal combination is further assumed when the sensory noise of two cues does not significantly differ from predictions of optimality. Re-weighting is indicated by a reduction in disparity cue weights when noise in the disparity cue is increased. Optimal re-weighting is indicated by empirical reductions in the disparity weight in line with optimal predictions. Sensitivity to incongruence is shown by an increase in sensory noise when cues are presented incongruently (inversed learnt mapping). ***Bottom***: The experiment was split into four sessions, with the key markers – combination, re-weighting, and incongruence sensitivity – measured across three consecutive sessions, preceded by a training session.

*Combination* was assessed via precision improvements or, in other words, a reduction of sensory noise in the combined cue conditions (e.g. Disparity + Size, DS) compared to the best individual cue condition (e.g. Disparity D, or Size S). The magnitude of precision improvement was further tested for deviations from optimal predictions. *Re-weighting* was assessed by measuring changes in relative cue weights for the disparity cue, before and after adding noise to the disparity stimulus. Empirical weights that indicate how much relative weight is placed on the disparity cue during integration was measured via cue-conflict-induced biases. Predicted weights that indicate how much relative weight the ideal observer would be expected to place on each cue, were measured via the relative sensory noise of the individual cue conditions. Re-weighting would be indicated by a reduction in disparity weight, with increasing noise in the disparity cue. *Sensitivity to mapping congruence* was assessed by measuring the reduction in precision, that is, the increase in sensory noise when the cues were presented in an incongruent, compared to congruent, mapping.

All three markers were measured across three separate sessions, with both cue pairings measured within the same sessions. Prior to this, participants completed a first session in which they underwent pre-assessment (section 2.6.1) and training based on associative learning (section 2.6.2). All sessions were conducted within the same week.

### 2.2. Participants

#### 2.2.1. Power and sample size estimation

For this experiment, we aimed to include 60 healthy adults in the analysis, based on simulation-based power estimations (Scarfe, 2022; Scheller & Nardini, 2023) to detect a significant cue combination effect (precision benefits) at group-level as well as resource availability. The power to reliably detect a cue combination effect depends on the relation between measurement noise and the maximally possible effect size. The latter, in turn, depends on participant-specific characteristics, such as the absolute sensory noise of the individual cues as well as their cue noise ratio (see also Scheller & Nardini, 2023). These sample characteristics cannot be predicted in advance; however, we used pilot testing to determine a range of likely participant-specific characteristics that informed our power simulations about possible sample parameters. For more details on the power analysis and visualized power curves, see supplementary material.

### 2.2.2. Initial sample & exclusion criteria

In total, 105 participants were recruited for the experiment. Out of these, 27 failed the stereo acuity pre-assessment (see below) and did not proceed to the training nor testing sessions. Out of the 78 participants that completed the tasks, data from two participants were excluded due to high lapse rates (> 10% of trials, across sessions). Furthermore, data from individual participants was excluded on a marker- and cue-pair basis, if their sensory noise values exceeded 2.5 times the IQRs of the group-level noise distributions.

#### 2.2.3. Final sample

The final sample comprised of 78 participants (55 female; Age: 22.6±4.3 years), 70 of whom were included in the familiar-familiar cue combination analysis (50 female; Age: 22.7±4.3 years), and 63 of whom were included in the familiar-novel cue combination analysis (46 female; Age: 22.5±4.3 years). In total, the cue combination analysis included complete data across both familiar-familiar and familiar-novel cue pairings from 58 participants (43 female; Age: 22.6±4.4 years).

### 2.3. Conditions

Conditions were defined based on cue availability, with all participants completing unimodal and bimodal sensory conditions of each of the two cue pairings (familiar-familiar, familiar-novel). Within the unimodal conditions, participants completed the depth discrimination judgement task using the three primary sensory cues alone: Disparity only (D), Size only (S), and Audio only (A). In the bimodal conditions, participants completed the task using either a congruently paired cues, incongruently paired cues, or congruently paired but conflicting cues.

#### 2.3.1. Single cues/unimodal

The disparity-only condition (D) utilized stereo vision to simulate object depth by introducing small horizontal displacements in dot locations, creating retinal disparity. The size-only condition (S) introduced object depth through size-distance constancy, where objects further away project smaller retinal images. When unimodal visual cues were presented, the other visual cue was held constant. In the audio-only condition (A), the central square was not visible, but the novel audio cue that provided information about the location of the object in depth was presented though headphones.

#### 2.3.2. Cue pairings/bimodal

To measure cue combination of familiar cues, the disparity cue changed in depth together with the size cue in a natural, correlated fashion (DS). To measure cue combination of familiar and novel cues, the disparity cue changed in depth together with the audio cue in the newly-learnt, correlated fashion (DA). To test for re-weighting, the uncertainty of the disparity cue was increased (Du) by introducing a variable jitter in the offset between the corresponding points in the left and right images. Furthermore, conflicting bimodal conditions were included. That is, a constant conflict was introduced between the disparity and the second, paired cue in the reference stimulus. Here, the disparity cue remained at reference level, while the size or audio cue was presented at a fixed offset of 1.5 times the Just Noticeable Difference (JNDs) between the cues. To avoid cross-modal adaptation, presentation of this offset was presented in positive and negative directions (± δ), leading to the four conditions DS_conflict_+, DS_conflict_-, DA_conflict_+, and DA_conflict_-. Positive and negative conflicts were presented in a randomized, counterbalanced fashion. The comparison stimuli did not include any conflicts between the cues. To test for sensitivity to mapping incongruence, bimodal conditions were included that contained an inverse mapping to the natural (familiar-familiar, DSi) or learnt (familiar-novel, DAi) mapping in the comparison stimulus. All sensory cue conditions were presented in a blocked fashion in a random order to reduce the influence of learning effects and avoid confusion through frequent switching of cue conditions.

### 2.4. Stimuli

#### 2.4.1. Visual cues

The visual stimulus comprised of a random dot stereogram (RDS), with dense (20 dots/ deg^2^) black and white dots arranged in a square shape (6° x 6° visual angle at screen level). This square was the target and was moved along the depth plane via the different sensory cues (binocular disparity, size, audio). The square was placed within the cut-out (7.5° x 7.5° visual angle) of a rectangular background shape (12° x 16° visual angle), which was used to prove a stable reference at screen level and to promote a stable vergence posture. The target square and background rectangle were made up of a dense display of individual dots (0.15° in diameter), for which exact placement (placed on an equally spaced grid with added random noise) and colour (black, white) were drawn randomly on each trial in order to reduce effects of visual adaptation. A white fixation cross was presented within a 1° square hole in the centre of the target square at screen depth. The visual stimulus was presented on a 23” monitor (ThinkVision, Lenovo), placed at 138cm from the participant. Viewing distance was confirmed at the beginning of each session.

The square moved in depth via the two different sensory cues: binocular disparity and size, which were mapped onto depth via their veridical, physical relationship. In other words, the magnitude of change induced in each cue to simulate a change in depth of 1cm corresponded to the veridical/natural magnitude of change in the physical features. Stereo images were presented via Psychtoolbox’s anaglyph stereo function and a pair of red-blue anaglyph glasses. A change in binocular disparity, simulated via a horizontal displacement of the stereo images in the left and right eye was given by:

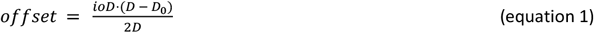

Where *ioD* is the participant’s interocular distance, *D*_*0*_ is the screen distance, and *D* is the normalized target distance. To measure re-weighting, the offset between the left and right images was varied randomly for each individual dot (*SD* = 27 pixels), reducing the reliability of the disparity cue. Note that, while this introduced random noise to the disparity cue, the average of all dots remained fixed to the chosen depth.

A change in depth via the size cue was induced via

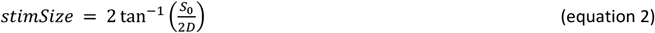

Where *S*_*0*_ refers to both the width of the square and the diameter of the individual dots at screen level, in centimetres. The size of the dots was altered alongside the size of the square to prevent changes in density, which could function as an additional cue if not controlled for.

#### 2.4.2. Auditory cue

The novel auditory cues that were mapped to depth consisted of pure tones with varying frequencies, played for the duration of the visual presentation (1s) at 48kHz sampling frequency, with a 10ms raised cosine on- and offset ramps. Screen depth was mapped to a frequency of 600 Hz. A simulated 1 cm change in the depth domain corresponded to a 0.015 semitone (1.5 cents) change in pitch. As pitch sensitivity of the human ear varies with frequency, all tones were loudness-adjusted such that the tone’s acoustic loudness, determined via the ISO 532-1 (Zwicker) method, was equal across all stimuli. This was done to the avoid use of perceived loudness as an additional cue.

Crucially, to avoid tapping into possible, pre-existing cross-modal correspondences (Jiang et al., 2024; Parise & Spence, 2009), the depth to pitch mapping was randomly assigned and counterbalanced across participants. Of the included participants, 36 learnt a mapping where pitch increased with increasing distance, while 42 of the participants learnt a mapping where pitch decreased with increasing distance.

#### 2.4.3. Stimulus depth ranges

Stimulus depth ranges and increment sizes that were used in testing were determined in a set of pilot experiments. As visually crossing the plane of the screen (at which binocular disparity equals zero, if fixating the screen) could provide a strong additional cue to relative depth, the depth plane was split into two ranges: one in front and one behind the screen. Individual participants were assigned to either of the ranges, based on their performance on the stereo-vision pre-assessment. Of the included participants, 41 were presented with stimuli that appeared to pop out in front of the computer screen (1-21cm closer; reference 11cm closer than screen; see Figure 2), while 36 were presented with stimuli that appeared to pop in to the computer screen (1-21cm further; reference 11cm further than screen). The depth range was normalized, with the reference being 0, and divided into 14 equal steps between +/–1 and +/-0.1 in increments of 0.15. Hence, the possible stimulus depths were at +/-1cm, 2.5cm, 4cm, 5.5cm, 7cm, 8.5cm, 10cm from the reference.

**Figure 2:**
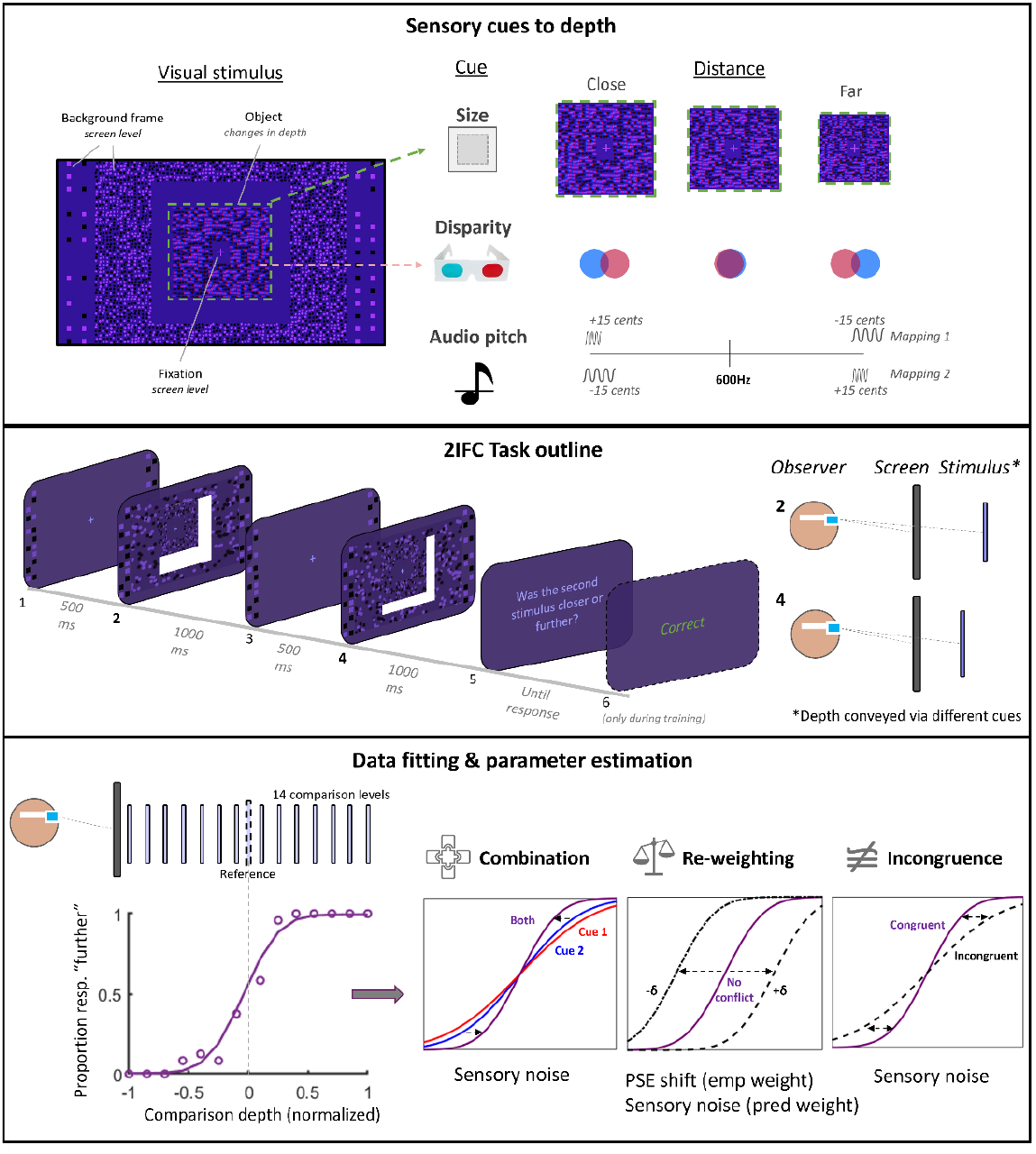
Top: Stimuli comprised visual (size, disparity; familiar) and auditory (pitch; novel) cues to depth. The familiar cues followed a natural mapping of depth, based on the physics of real-world cue behaviour. With increasing distance, the object would become smaller (size cue). The offset between left and right image would increase with increasing distance from the screen, with the directionality of the offset indicating the stimulus being closer or further than the screen (disparity cue). The novel audio cue was mapped to depth with either increasing pitch or decreasing pitch with further distance. The mapping direction was counterbalanced across participants. ***Middle***: Temporal outline of the 2IFC task. Two stimuli were presented after each other at different levels of depth, indicated in the sketch on the right, with a 500ms ISI. Depth was conveyed via individual cues or via two cues at the same time. ***Bottom***: Sensory noise parameters were estimated by fitting psychometric functions (PF) to the proportion of trials on which participants responded that a specific comparison stimulus (different levels represented in pale purple in the sketch above) was further than the reference stimulus (presented with a dashed outline in the sketch above, aligned with 0 of the PF). To assess combination, we estimated the sensory noise of the single and combined cue conditions from changes in the PF slope (equation 5). To assess re-weighting, we estimated the absolute mean shift in PSEs of the conflict condition PFs, relative to the no conflict condition. Weights for the disparity cue were derived from these shifts relative to the offset (*δ*; equation 7). Also, sensory noise was estimated for the single cue conditions to assess the relative predicted weight for the disparity cue (equation 8). To assess incongruence sensitivity, we estimated the sensory noise of the combined congruent and combined incongruent conditions.

### 2.5. Task

The depth discrimination task employed a two-interval forced-choice (2IFC, Figure 2) paradigm, whereby two stimuli were presented in succession via one or two cues, depending on the specific sensory cue condition. On one of the two presentation intervals, the stimulus was presented at a fixed distance (reference stimulus), while on the other presentation interval, the stimulus was presented at one of 14 different distance levels (comparison stimulus), interspersed across the depth range. On each trial, the order in which reference and comparison stimulus appeared was randomly selected. The order in which the different comparison depth levels were presented was randomized. The method of constants was used for stimulus selection with 12 repetitions per stimulus level and sensory condition.

After showing the two stimuli in succession, participants had to decide whether the second stimulus was closer or further than the first stimulus, by pressing a button on a handheld controller. Response feedback was only given during training, but not during testing sessions. To prevent long stretches of non-discriminable trials, 4% of “easy” trials were introduced where the most extreme stimulus distances were presented in the same trial. Participants received an hourly base rate payment of £5. To enhance engagement and maintain attention, participants further obtained performance-dependent winnings and performance feedback after every 50 trials. The maximal performance-dependent winnings per condition (176 trials) were £2, adding up to £14-£18 per session. On average, 58% of the final payment was based on performance-dependent winnings.

### 2.6. Experimental session structure

The first session consisted of a pre-assessment task and training. During the training, trial-based feedback was provided. The second, third and fourth sessions included all testing conditions to determine cue combination (session 2), cue re-weighting (session 2 and 3), and sensitivity to incongruence (session 4).

#### 2.6.1. Pre-assessment Task

Before the main tasks, participants were assigned to one of two groups using different stimulus depth ranges. To ensure participants can perceive depth from disparity at the selected ranges, a pre-assessment task presented 20 disparity-only stimuli, ten at the closest and ten at the furthest points of the range, as well as five stimuli at the same level as the screen. Participants had to judge whether stimuli appeared to pop out, pop in, or neither. If participants responded incorrectly on at least 20% of the close stimuli, they were assigned to the group using the far stimulus range, or vice versa. If they responded incorrectly on at least 20% of the close and far stimuli, or on the flat stimuli, their stereo acuity was deemed too low to participate in the experiment. Overall, around half of the participants (50.8%) were assigned to the closer stimulus range, while the other half was assigned to the further stimulus range.

#### 2.6.2. Training

During training, participants conducted 2IFC depth discrimination judgements in familiar unimodal conditions (D, S) and combined conditions (DS, DA) with 300 trials per condition. The novel auditory-only condition (A) was excluded prior to training as pitch and depth mapping were established during this session through exposure to the coupling of disparity and audio cues (DA). In contrast to the testing sessions, participants received feedback after each trial to enforce mapping learning between the familiar disparity and the novel audio cues. The condition order was randomized but remained consistent for each participant across different days to minimize order effects.

#### 2.6.3. Main testing task

In the main testing sessions (sessions 2-4), participants conducted all unimodal conditions (sessions 2 and 4: D, S, A; session 3: Du, S, A) and bimodal congruent conditions (sessions 2 and 4: DS, DA; Session 3: DuS, DuA) with 150 trials per condition. They received no feedback after individual trials. To maintain attention and engagement, they received accuracy feedback for each block of 50 trials, as well as performance-dependent winnings. The audio-only condition was presented after the disparity-audio condition (DA) to ensure that participants were recently exposed to the mapping. Conditions assessing re-weighting (session 2: DS+, DS-, DA+, DA-; session 3: DuS+, DuS-, DuA+, DuA-) or sensitivity to incongruence (session 4: DSi, DAi) were presented after those testing for combination (D, Du, S, A, DS, DA, DuS, DuA) to avoid potential residual effects that could result from continuously presenting cue conflict (e.g. adaptation) or incongruent stimuli (e.g. unlearning the mapping). Precautions against these effects were already implemented in the tasks (intermixing of conflict offset directionality, enhanced presentation frequency of congruent stimuli), however, they are impossible to fully exclude.

### 2.7. Parameter estimation

All three markers of native perception were investigated by comparing different perception-dependent psychometric parameters (see Analysis and Figure 2, bottom). Combination and incongruence sensitivity were based on the analysis of *sensory noise* (σ), while re-weighting was based on shifts in the *point of subjective equivalence* (PSE) in the conflict conditions (to derive empirical weights, *w*^*emp*^) as well as the sensory noise of the single cue conditions (to derive predicted weights, *w*^*pred*^). To estimate these parameters, we fitted psychometric functions to the response data of the 2IFC task. The psychometric function for a 2IFC discrimination task with a reference at zero can be formalized as:

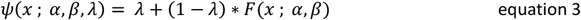

where *ψ* describes the proportion of responses ‘further’ as a function of the stimulus depth *x*, while *λ* indicates the lapse rate and determines the upper and lower asymptote of the function. The function *F*(*x* ; *α, β*) is modelled by a cumulative normal distribution:

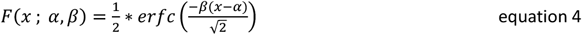

where *α* describes the midpoint of the function and directly indicates the PSE. Sensory noise links to the normal distribution’s standard deviation, which is the inverse of the slope of the function, described by *β*.

Psychometric functions were fitted to single cue conditions using Bayesian criterion fits with uniform priors, implemented in Palamedes (version 1.11.9; Kingdom & Prins, 2016). Posterior distributions were sampled via MCMC, implemented in JAGS (version 4.3.1; (Plummer, 2012). Parameter estimation was optimized to reduce imprecision in the parameters of interest for the different markers: When estimating sensory noise parameters, *α* was fixed to zero as deviations from zero would be highly improbable and theoretically not meaningful. In cue conflict conditions, in which a shift in PSE is theoretically interpretable and expected, *α* was allowed to vary between -1 and 1, demarcating the minimum and maximum of the full stimulus range. Realistically, however, values were expected to range between zero and the cue conflict magnitude (*δ*). The slope parameter, *β*, that was linked to the cue’s sensory noise and that provided a key value of interest, was not constrained further than the default (which allowed positive values for the slope only). The lapse rate parameter *λ* was constrained between 0 and 0.2, allowing to estimate individual lapse rates up to 20%. This was theoretically motivated: While lapses on more than 10% of trials would indicate very poor data quality, limiting estimation to lower lapse rates would enhance the possibility of including poor data in the final analysis. Instead, we decided to exclude data from participants whose lapse rates were estimated to be higher or equal to 10% across sessions. The *λ* search range was initialized at 0.02.

To reduce measurement noise and the chance of conflating lapses with sensory noise (see Scheller & Nardini, 2023, Supplementary Material), an initial, hierarchical, Bayesian model was fitted for each participant and session to estimate *λ* across sensory conditions. This draws on the advantage of using more data to enhance estimation precision in the lapse parameter (Prins, 2024). Lapses are not assumed to be dependent on each sensory condition, but rather differ for each participant and session. By subsequently fixing *λ* to the individual and session-specific values, we were able to reduce bias and imprecision in the estimation of our parameter of interest, *β*, in the cue-specific conditions.

### 2.8. Calculation of sensory noises and weights

As sensory noise (σ) of the discrimination performance results from the comparison between two stimuli, it can be derived from the variance of the underlying Gaussian 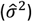, or the function’s slope (*β*) via:

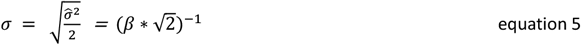

To scale the cross-modal cue conflict between disparity and size, or disparity and audio (to measure re-weighting), we further estimated the Just-Noticeable Difference (JND) from sensory noise via:

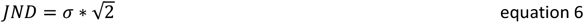

The JND is a useful psychophysical unit describing the difference between two stimuli that can be just noticed and, hence, discriminated, by a participant. The JND was used to scale the difference in depth in the cue conflict conditions (*δ*), which allowed to derive empirical cue weights (*w*^*emp*^). As weights are derived from estimated parameters, which carry inherent measurement noise, the conflict between two cues needs to be large enough to allow for a testable difference between which cues are being followed. At the same time, cue conflicts need to be small enough to not induce an obvious discrepancy, as strongly noticeable discrepancies would reduce the probability with which participants integrate the cues (Gepshtein et al., 2005; Körding et al., 2007). Following best practice recommendations (Rohde et al., 2016), the cue conflict was set at ± 1.5 JNDs for the size or auditory cue for each cue pairing.

Empirical cue weights for the non-noisy 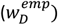 and noisy 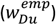 disparity cues were derived from the conflict-induced shift in PSEs (*μ*_*PSE*_), averaged for positive and negative conflicts, relative to the maximum conflict:

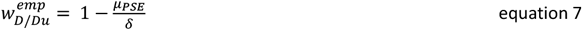

According to the optimal cue combination model, cues are weighted by their individual reliabilities. As such, one would expect that empirical cue weights align with model predictions. Predicted cue weights for the non-noisy 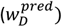 and noisy 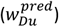 disparity cues were derived from the relative sensory noise of the individual cues via:

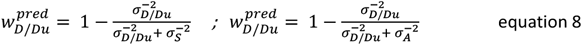

Furthermore, to quantify the differences between cue pairings, we calculated indices for combination, re-weighting, and incongruence sensitivity for each participant. These indices depended on their definition of the process, that is, combination is evidenced by a reduction in sensory noise relative to the best individual cue, i.e.:

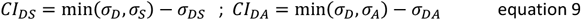

A positive *CI* indicates an enhanced advantage relative to the best individual cue. In the preregistration, we suggested scaling this relative to the benefit that can be obtained based on optimal predictions, however, large inter-individual variability and variable cue noise ratios limit the comparability of this index in a group-based analysis. Hence, the CI indicated the absolute benefit obtained from two cues.

As re-weighting is evidenced by a reduction in cue weights when sensory uncertainty is added to the cue, the re-weighting index for each cue pairing can be determined as:

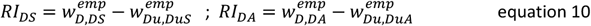

A positive *RI* indicates that more weight is placed on the disparity cue when it is not noisy, compared to when it is noisy.

Sensitivity to incongruence is evidenced by a reduction in precision, i.e., enhanced sensory noise, when the established cue mapping is reversed. For instance, when the cue mapping is frequently experienced and strongly established, such as between highly familiar cues, the presentation of an anti-correlated cue mapping would result in an enhanced sensory noise. This can result either through enhanced fusion (Ernst, 2007) or increased separation and switching between cues (Nardini et al., 2010). Hence, the incongruence sensitivity index *IS* is calculated via the difference in sensory noise between the congruent and incongruent conditions:

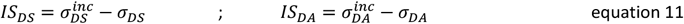

An IS that deviates strongly from zero indicates an enhanced sensitivity to incongruence. More positive values mean that incongruence leads to enhanced sensory noise, which can result from fusion or switching between the cues, while more negative values indicate a reduction in sensory noise as a result of incongruence, which may indicate that a single cue was used most of the time.

### 2.9. Analysis

#### 2.9.1. Cue combination (Q1)

Combination predicts that precision of the bimodal estimate will increase (I.e., sensory noise σ will be reduced), relative to the best unimodal estimate. That is:

> To test **H1a**, we will ask if σ_*DS*_ < *min*(σ_*D*_, σ_*S*_) using two-tailed Wilcoxon Signed-Rank Tests.
>
> To test **H1c** ask if σ_*DA*_ < *min*(σ_*D*_, σ_*A*_) using two-tailed Wilcoxon Signed-Rank Tests.

Furthermore, statistically optimal combination following maximum likelihood estimation (equation 12) predicts that the sensory noise of the bimodal estimate does not deviate from model predictions, hence:

> To test **H1b**, we will ask if σ_*DS*_ differs from 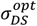 using two-tailed Wilcoxon Signed-Rank Tests.
>
> To test **H1d** ask if σ_*DA*_ differs from 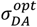 using a two-tailed Wilcoxon Signed-Rank Tests.

The optimal integration model describes how sensory noise in the combined conditions decreases in accordance with the individual cue variances. Hence, predictions for optimal combination following the maximum-likelihood model are given by:

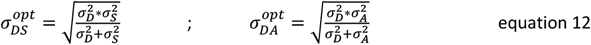

To ascertain that the possible absence of cue combination effects was not purely resulting from large differences in cue variances and, hence, small possible effect sizes (see Scheller & Nardini, 2023 for more info), we tested for combination in a sub-sample of participants. This sub-sample only included individuals whose individual cue sensory noise ratios were below 1.5. As indicated in our power simulations, the ability to detect significant cue combination effects strongly decreases with larger cue variances.

#### 2.9.2. Re-weighting (Q2)

Cue re-weighting predicts that more weight will be given to the more reliable sensory modality. As sensory noise increased in the disparity stimulus in session 3, relative to session 2, we would expect lower disparity cue weights when noise is added (i.e., DuS and DuA) compared to when the disparity cue is not noisy (i.e., DS and DA), that is:

> To test **H2a**, we will ask if 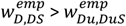 using one-tailed Wilcoxon Signed-Rank Tests.
>
> To test **H2c**, we will ask if 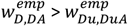 using one-tailed Wilcoxon Signed-Rank Tests.

Additionally, the predicted weights 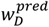 and 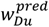 describe how much the disparity cue should influence the final percept if combination takes the relative reliabilities of the individual cues into account. As such, the reduction in empirically measured disparity weights will be compared with the reduction that would be expected based on reliability-weighted, optimal cue combination. If re-weighting follows optimal predictions, we expect no significant differences in predicted and empirically determined weight reductions:

> To test **H2b** we will ask if 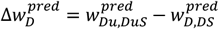 does not differ from 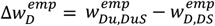, using two-tailed Wilcoxon Signed-Rank Tests.
>
> To test **H2d** we will ask if 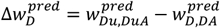 does not differ from 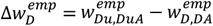, using two-tailed Wilcoxon Signed-Rank Tests.

#### 2.9.3. Sensitivity to incongruence (Q3)

Sensitivity to incongruence would predict that precision of the incongruent bimodal estimate reduces (i.e., sensory noise increases) relative to the combined congruent cue. To determine which of the two cues participants followed mostly, we fitted participants’ responses against depth changes in either of the two cues and determined the better fit.

> To test for incongruence sensitivity of the familiar cue pair (**H3a**), we asked whether 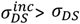, that is, whether the sensory noise of the incongruent condition is larger than the sensory noise of the congruent condition, using two-tailed Wilcoxon Signed-Rank tests.
>
> To test for incongruence sensitivity of the familiar-novel cue pair (**H3b**), we asked whether 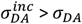,using two-tailed Wilcoxon Signed-Rank tests.

#### 2.9.4. Differences between cue pairings (*H4*)

For each marker and participant, we further calculated indices for cue combination (*CI*_*DS*_, *CI*_*DA*_), re-weighting (*RI*_*DS*_, *RI*_*DA*_), and incongruence sensitivity (*IS*_*DS*_, *IS*_*DA*_), that quantified the degree with which these different markers were expressed. Positive values indicate that participants perceptually benefitted from combining the cues (*CI*), reduced the weight of the disparity cue (*RI*) when noise was added, relative single cue weights, and showed sensitivity to incongruence (*IS*). To test whether novel-familiar cue combination is processed similarly to familiar-familiar cue combination on all these markers, we contrasted the indices between the cue pairings. To that end, we first tested whether combination, re-weighting, and incongruence sensitivity indices were larger for familiar-familiar than familiar-novel pairings at the group-level:

> That is, to assess **H4** we asked if *CI*_*DS*_ > *CI*_*DA*_, if *RI*_*DS*_ >*RCI*_*DA*_, and if *IS*_*DS*_ > *IS*_*DA*_, using two-tailed Wilcoxon Signed-Rank Tests.

Secondly, we determined how many participants showed combination, re-weighting, and incongruence sensitivity effects in the expected direction for the familiar-familiar pairing versus how many participants showed such effects for the familiar-novel pairing.

Here we asked if the number of participants that showed *CI*_*DS*_ > 0 differs from the number of participants that showed *CI*_*DA*_ > 0, whether the number of participants that showed *RI*_*DS*_ > 0 differs from those that showed *RCI*_*DA*_ > 0; and finally, whether the number of participants that showed *IS*_*DS*_ > 0 differs from the number that show *IS*_*DA*_ > 0, using McNemar’s test for paired samples.

### 2.10. Exploratory analyses

#### 2.10.1. Intra-individual stability and inter-individual differences

Importantly, sensory noise measures may be impacted by variability resulting from measurement noise or a general instability of cue combination over time. To the best of our knowledge, no previous study has shown whether cue combination can be reliably and repeatedly estimated across within the same individuals across time. The large sample and repeated testing structure of this experiment therefore provided a unique opportunity to estimate repeatability for cue combination effects within the same individuals. This is particularly useful for inferring whether variability in the whole sample can be attributed to sources of within- or between-individual factors. Within-individual factors may result from measurement noise, which affects every single measurement time point. It may further result from inconsistency of cue combination within participants across days, resulting from extraneous factors such as lack of attention, strong learning effects, or switching of information processing strategies. On the other hand, variability in cue combination effects in the sample may be the result of between-individual factors. This would suggest that inter-individual differences exist in the ability to combine sensory cues (optimally).

To assess whether cue combination variability in the group resulted from intra-individual variation, we estimated the repeatability of sensory noise measurements over sessions 2 and 4, which included the same sensory conditions. Mixed effect models, which allow to partition subject- and group-based variances on different hierarchical levels, were used to quantify repeatability. Within those models, repeatability was estimated in accordance with (Stoffel et al., 2017) via the variance among group means 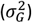 over the sum of group- and data-level (residual) variance 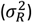.

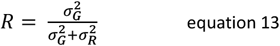

Reduced repeatability indicates high variability in cue combination within the same individual. On the other hand, highly repeatable measurements, indicated by larger repeatability indices, would suggest that cue combination effects are rather stable within the same individual. Estimate uncertainty, assumed Gaussian, was quantified by means of parametric bootstrapping, using 10000 samples. Significance of repeatability estimates was implemented by likelihood ratio tests (see Stoffel et al., 2017).

Due to the repeated exposure to coupled cues in the DA condition, it was still possible that cue combination performance changed systematically across the experiment. As such, we assessed whether group-level cue combination effects differed between day 2 and day 4.

Furthermore, we conducted intra-individual correlations to assess whether individual differences in one marker may explain differences in the other markers. To that end, we cross-correlated the functional indices (CI, RI, IS) within each cue pairing.

## 3. Results

### 3.1. Excluded data

Some participants showed particularly high sensory noise values in individual cue conditions, suggesting they had difficulty discriminating stimuli based on those cues. We therefore excluded data from participants whose sensory noise values in the single cue conditions, or cue weights, were more than 2.5 times IQR from the median of the group. Two participants showed very high sensory noise levels in the disparity cue (present in familiar-familiar, and familiar-novel cue pairings) across sessions and were therefore excluded from all analyses. A further two participants showed high disparity noise only in the last session. Their data were therefore excluded from fusion analysis. Based on sensory noise in the size cue, data from three participants had to be excluded to assess markers of the familiar-familiar cue pairings. Similarly, some participants did either not learn the cue mapping correctly or showed very low precision in pitch discrimination and were excluded based on the criteria outlined above. This led to the exclusion of data from 12 participants when testing for familiar-novel cue pairings.

### 3.2. Pre-registered analyses

#### 3.2.1. Combination

Combination predicts that sensory noise σ of the bimodal estimate will be reduced relative to the best unimodal estimate. Figure 3 shows sensory noise across conditions, indicating a reduction in sensory noise in the bimodal estimate, compared to the best individual cue, for the familiar cues (size and disparity; Figure 3, top left). A two-tailed Wilcoxon Signed-Rank Test indicated that this reduction was significant (H1a: *V* = 1888, *p* < .001). Furthermore, performance in the bimodal condition did not differ significantly from optimal predictions (H1b: *V* = 1052, *p* = .266).

**Figure 3:**
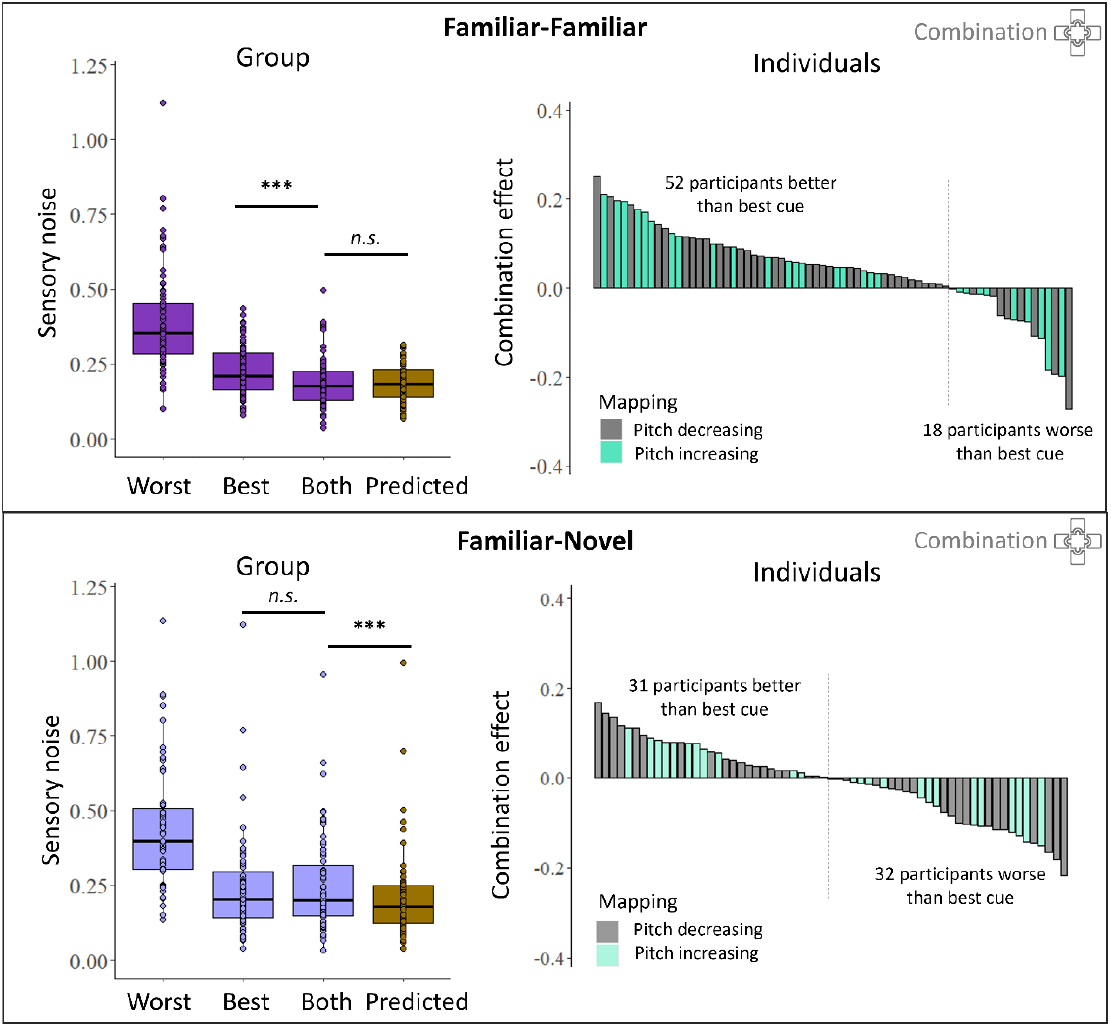
Combination results for familiar-familiar (disparity-size, *top panels*) and familiar-novel (disparity-audio, *bottom panels*) cue pairs. *Left panels*: Box plot shows sensory noise distribution for the individually-determined worst and best single cues, and for the bimodally presented cues in purple. Predictions that follow the optimal integration model are shown in dark yellow. *** *p* < .001; *n.s*. not significant. *Right panels*: Distribution of combination effects in the sample, given by the difference in sensory noise of the bimodal condition relative to the best single cue condition. Every bar represents one participant. Combination effects are ordered by size for visualization purposes to highlight the proportion and variability of combination in the sample. The participant-specific depth-to-pitch mapping is shown in different coloured bars, showing that the ability to combine did not depend on the novel cue mapping that was learnt.

On the contrary, when using the novel audio cue in combination with the familiar disparity cue, sensory noise was not significantly reduced relative to the best unimodal estimate (H1c: *V* = 920.5, *p* = .551; Figure 3, bottom left), suggesting that cues were not combined. Furthermore, sensory noise in the combined condition differed significantly from optimal predictions (H1d: V = 1456, *p* = .002), showing that the lack of combination was not due to an inability to detect small, optimal combination effects from the best cue.

Combination effects, expressed as the increase in precision (decrease in sensory noise) when two cues were presented together, relative to the best single cue, showed considerable variability between participants (Figure 3, right). When familiar cues were presented together, the majority of participants showed increases in precision relative to the best cue (52 out of 70; 74%), suggesting that they integrated the two cues. The other quarter of participants did not benefit from having both cues available, or even performed worse than with the best single cue (18 out of 70; 26%). This would suggest that they employed a different perceptual-decisional strategy, such as cue switching. When the familiar and novel cues were presented together, only about half of the participants showed positive precision increases (31 out of 63; 49%), while the other half did not benefit from having both cues presented together (32 out of 63; 51%). The degree of combination benefit did not depend on the participant-specific audio-disparity cue mapping direction, as can be seen by the distribution of grey and turquoise bars in Figure 3 (right panels).

As the noise ratio of the individual single cues strongly affects the ability to measure cue combination benefits (Scheller & Nardini, 2023), we ran several pilot studies to ensure that the chosen cue ranges were, on average, well matched across individuals. There was, however, residual variability in cue noise ratios in the included sample. The median cue noise ratio for the familiar cue pair was 1.57 (range: [1.01 5.03]), while the median cue noise ratio for the familiar-novel cue pair was 1.98 (range: [1.01 30.4]). A ratio of 1 would indicate that both cues were perfectly matched and would provide the best chances of detecting true combination (Scheller & Nardini, 2023). To ensure that the absence of combination was not merely the result of higher cue noise ratios. we further tested for combination benefits with the novel cue in a subset of observers (*N* = 17) that showed a cue noise ratio below 1.5. This analysis confirmed the group-level findings reported above: familiar-novel cue pairs were not combined at the group level, with the sensory noise of the bimodal cue being higher than optimal (*V* = 142, *p* < .001) and not different from the best single cue (*V* = 66, *p* = .644). Again, combination ability differed within the sample, with 8 observers showing a combination benefit and 9 observers showing no combination benefit.

#### 3.2.2. Re-weighting

By introducing a small, consistent conflict between the simultaneously presented cues, we measured the weight that was placed onto each cue during combination. Reliability-based cue re-weighting predicts that more weight should be given to the more reliable sensory modality Figure 4 shows participants’ weights for disparity in normal vs noisy conditions (*left*), comparing individuals’ degree of shift across conditions with predictions (*middle*), and showing the distribution of shifts across individuals (*right*). On a group level (Figure 4, *left*), when sensory noise in the disparity cue increased, the relative weight of the disparity cue reduced instantly, both when it was paired with the familiar size cue (H2a: *V* = 25, *p* < .001; *n* = 60) and when it was paired with the novel audio cue (H2c: *V* = 37, *p* < .001; *n* = 55).

**Figure 4:**
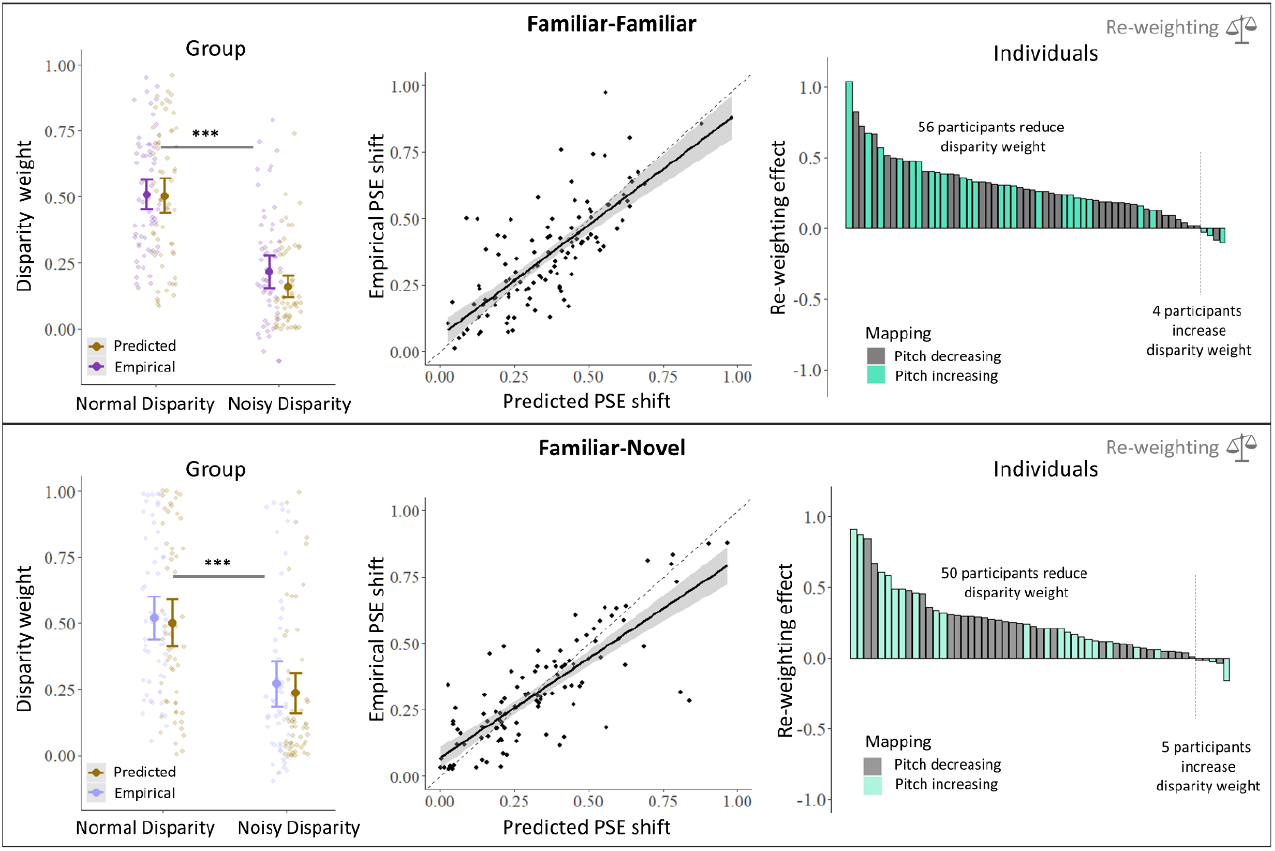
Re-weighting results for familiar-familiar (disparity-size, *top panels*) and familiar-novel (disparity-audio, *bottom panels*) cue pairs. *Left panels*: Mean empirical weights 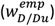 and predicted weights 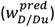 for the disparity cue as a function of noise in the stimulus disparity cue. Empirical cues were derived from PSE shifts in combined cue conditions (equation 7) while predicted weights were derived from sensory noise in the single cue conditions (equation 8). Error bars show 95% confidence intervals and single data points show the weights of individual participants. *** *p* < .001 *Middle panels*: The empirical shift in PSE of individual participants is plotted against the predicted shift in PSE, based on single cue uncertainty. Dashed black line indicates identity line. Bold black line and shaded area indicate a linear fitted regression model with standard error bands. *Right panels*: Distribution of re-weighting effects in the sample, given by the difference in empirical disparity cue weights between normal and noisy disparity conditions. Every bar represents one participant. Re-weighting effect is ordered by size for visualization purposes to highlight the proportion and variability of re-weighting in the sample. The participant-specific depth-to-pitch mapping is shown in different coloured bars.

Additionally, we tested whether this weight reduction differed from predictions of optimal integration. These comparisons showed that empirically measured weights reductions did not differ from optimal predictions both when the disparity cue was paired with the familiar size cue (*V* = 577, *p* = .222) and when it was paired with the novel audio cue (*V* = 669, *p* = .684). Additional exploratory analyses showed that individuals’ the optimal weight significantly predicted their empirical weight in both cases (familiar-familiar: *β* = 0.825; *t* = 14.35; *p* < .001, *r* = 0.8; familiar-novel: *β* = 0.753; *t* = 14.76; *p* < .001, *r* = 0.82; Figure 4, *middle panels*).

Individual re-weighting effects, indicating the decrease of disparity weight in the noisy disparity condition, relative to the non-noisy disparity condition, showed that most participants adjusted disparity weights in line with the stimulus noise. That is, when noise was added to the disparity cue in the stimulus, 56/60 (93%) participants put more weight on the size cue during combination.

Similarly, when the familiar and novel cues were presented together, 50/55 (91%) participants showed an increase in auditory cue weight when disparity was more noisy. The degree of re-weighting did not depend on the participant-specific audio-disparity cue mapping direction (Figure 4, right panels).

#### 3.2.3. Incongruence sensitivity

Sensitivity to mapping congruence would predict that precision of the incongruent bimodal estimate would reduce (i.e., sensory noise increases) compared to the combined congruent cue. Indeed, both cue pairs showed substantial sensitivity to incongruence, with the sensory noise of the incongruent condition being higher than the sensory noise of the congruent condition (see Figure 5, left panels). This increase in sensory noise was significant for both the familiar cue pair, disparity and size (*V* = 45, *p* < .001) and the familiar-novel cue pair, disparity and audio (*V* = 141, *p* < .001).

**Figure 5:**
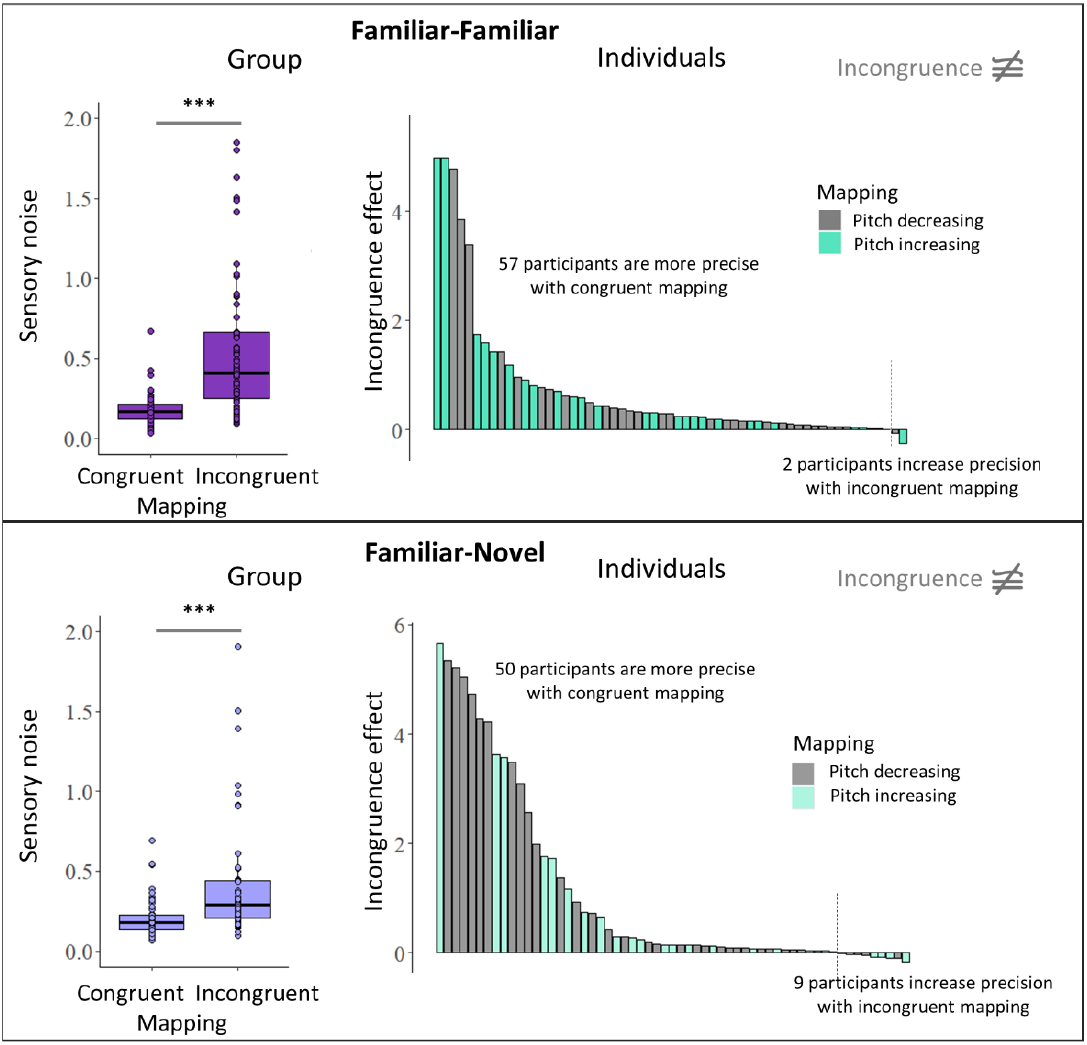
Incongruence sensitivity results for familiar-familiar (disparity-size, *top panels*) and familiar-novel (disparity-audio, *bottom panels*) cue pairs. *Left panels*: Box plot shows sensory noise distribution for the congruently paired and incongruently paired cues. For visualization purposes, values sensory noise values of 2 are not plotted here. *** *p* < .001. *Right panels*: Distribution of incongruence sensitivity effects in the sample, given by the difference in sensory noise of the bimodal incongruent condition relative to the bimodal congruent condition. Every bar represents one participant. Incongruence sensitivity effects are ordered by size for visualization purposes to highlight the proportion and variability in the sample. The participant-specific depth-to-pitch mapping is shown in different coloured bars.

Individual incongruence effects, indicating the decrease in precision/increase in sensory noise for the incongruent, relative to the congruent mapping condition, showed that precision decreased for the incongruent pairing in most participants and across cue pairs (Figure 5, right panels). Here, a reversed mapping of size and disparity lead to a decrease in precision in 57/59 (97%) participants. Similar, A reversed mapping of the novel audio cue and disparity led to a decrease in precision in 50/59 (85%) participants.

#### 3.2.4. Comparison of markers between cue pairs

To compare the degree of combination, re-weighting, and incongruence sensitivity between the two cue pairs, we contrasted the indices for each of these markers between familiar-familiar and familiar-novel cue pairs. Figure 6 plots these indices for each task. Positive values indicate that participants perceptually benefitted from combining the cues (*CI*), re-weighted the cues (*RI*) according to cue reliability, and showed sensitivity to incongruence (*IS*).

**Figure 6:**
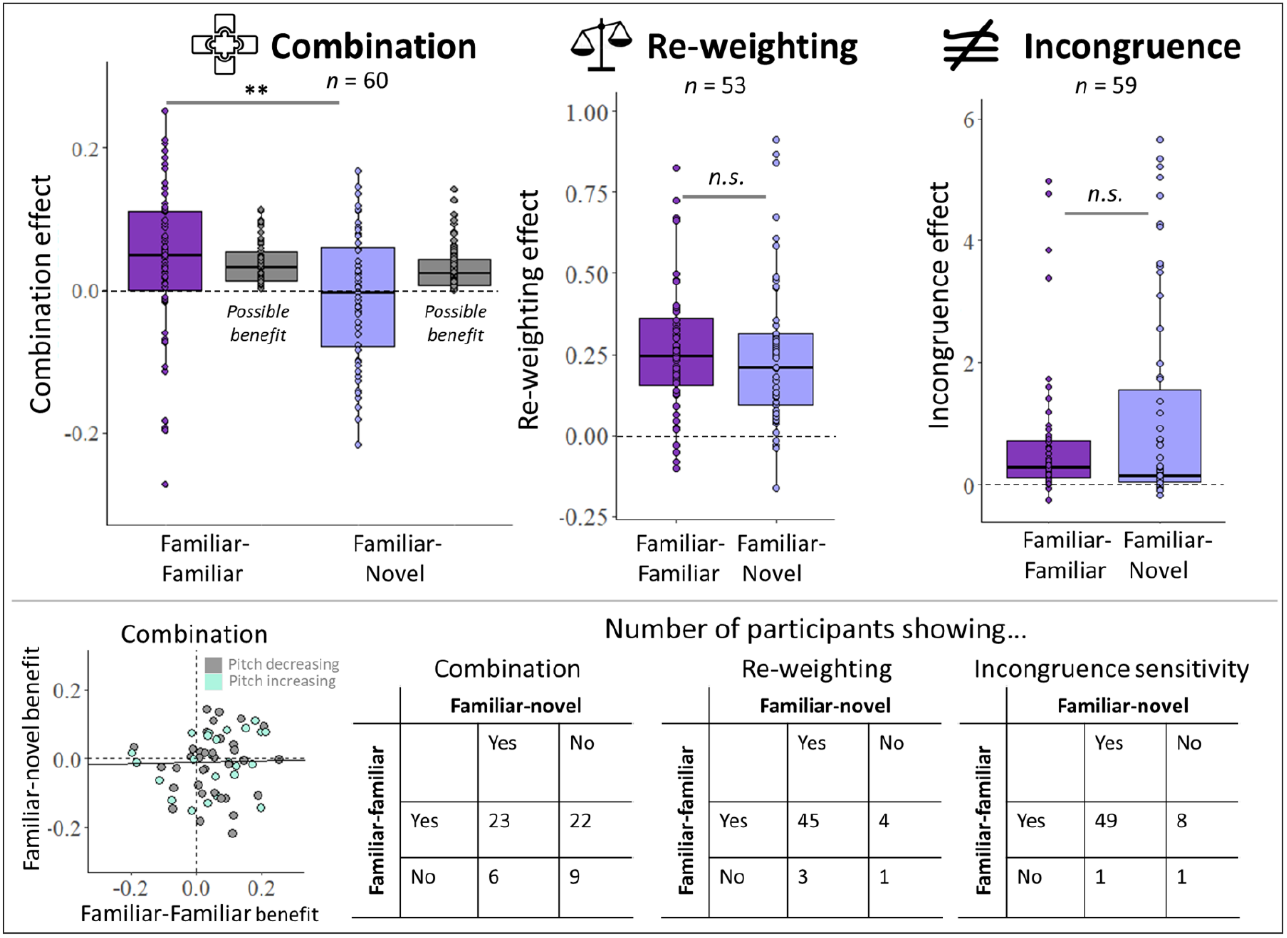
Combination, re-weighting, incongruence sensitivity indices contrasted between familiar-familiar (disparity-size) and familiar-novel (disparity-audio, *bottom panels*) cue pairs. The combination panel further shows the possible combination benefit, based on individuals’ optimal predictions, for reference. ** *p* < .01; *n.s. not significant*. Bottom panel: the left scatter plot shows the individual combination benefits for familiar-novel cues plotted against the individual combination benefits for familiar-familiar cues. Turquoise filled circles indicate those individuals who learned a positively correlated pitch mapping, grey circles indicate those individuals who learned a negatively correlated pitch mapping. Tables on the right show the number of participants that exhibited the three markers with either both cue pairs, one of the cue pairs, or none. A marker was classified as present if an individual’s index exceeded zero. See main text and analysis section 2.9.4. for more information.

Cue combination indices showed a larger benefit from combining the two familiar cues, compared to the familiar and novel cues (*V* = 1354, *p* < .001; Figure 6). Note that the possible benefit that people could obtain between the familiar-familiar and familiar-novel cue pairs did not differ significantly (*V* = 1115, *p* = .142), suggesting that the difference between the actual combination benefits was not due to differences in the ability to measure benefits. This reiterates the findings above showing clear, significant cue combination benefits at the group level in familiar-familiar cue pairings, and no significant group-level benefits for familiar-novel cue pairs. Re-weighting indices were not significantly different between the familiar-familiar and the familiar-novel cues (*V* = 724, *p* = .944; Figure 6). Lastly, incongruence sensitivity did not significantly differ between familiar-familiar and familiar-novel cues (*V* = 767, *p* = .375; Figure 6).

As shown above, there were strong individual differences in the degree with which each marker was expressed – especially in cue combination. Hence, we classified individual participants’ performance and compared the number of participants that showed each marker in both, either, or neither of the cue pairing, using McNemar test for paired factors. This showed a significant difference in the proportion of participants that combined familiar-familiar cues relative to familiar-novel cues (*χ*^*2*^ = 8.04, *p* = .005). Out of 60 participants, the majority combined both cue pairs (23/60, 38%) or only the familiar cue pair (22/60, 37%). On the other hand, participants that did not combine familiar cue pairs were also less likely to combine novel cues (9/60, 15%). Combining novel cue pairs, but not familiar ones was the least likely (6/60, 10%). Individual combination benefits were not significantly correlated between the familiar-familiar and familiar-novel cue pairs (*r* = 0.14; *p* = .28), suggesting that there was no cue-independent, individual proclivity to combine two cues to depth.

Contrasting the number of participants that dynamically re-weighted the cues showed that the majority of participants re-weighted cues in accordance with their reliabilities in both cue pairings (85%). There was no significant difference between cue pairs, suggesting that cues were not re-weighted more in a specific pairing (*χ*^*2*^ = 0, *p* = .999). Lastly, the majority of participants (83%) showed sensitivity to cue pairing incongruence in both cue pairs, however, it was significantly more prevalent in the familiar-familiar cue pair compared to the familiar-novel cue pair (14% vs 2%; *χ*^*2*^ = 4, *p* = .046).

### 3.3. Exploratory post-hoc analyses

#### 3.3.1. Intra-individual stability and inter-individual differences

While group-level results suggest that familiar-novel cues were not combined, there was considerable variability amongst participants. Importantly, this variability may result from different underlying noise sources: It can reflect *measurement noise*, inherent to data collection and fitting, *intra-individual variability*, such as state-dependent changes in alertness or attention that can vary across time, or it can reflect true, functional *inter-individual differences*. The contribution of the different underlying noise sources, however, is unclear. A key difference between them lies in their stability across multiple measurements: while measurement noise and intra-individual variability lead to changes in an observer’s estimates over time, true inter-individual variability would lead to more repeatable measures for the same observer over time. In the present study, we tested conditions that allowed to assess combination of the different cue pairs in two sessions (sessions 2 and 4). There was no additional training between the sessions. This repeated testing across two sessions allowed to determine the repeatability of estimates. In other words, it allowed to put the relative variability within participants (across sessions) in relation to the variability between participants (Nakagawa & Schielzeth, 2010). If a substantial part of the observed variability (Figure 3, *right panels*) was resulting from measurement noise or intra-individual variability, repeatability should be low. If, on the other hand, the observed variability arises from true differences between observers, the repeatability should be high.

Hence, measurement repeatability provides a principled way to assess whether large parts of the variability can be attributed to sources within individuals or between individuals. The repeatability index *R* was calculated based on mixed effect models as the group variance, divided by the summed group and residual variances (Stoffel et al., 2017, 2019). Observers were entered into the model as random effects. Uncertainty of R estimates was assessed via parametric bootstrapping (10000 times), and significance of repeatability was assessed via likelihood ratio tests and permutation of residuals (1000 times; see Stoffel et al., 2017).

The estimation of individual sensory cue noise was highly repeatable across unimodal (D: *R* = 0.516, *d*_*LRR*_ = 24.2, *p* < .001; S: *R* = 0.412, *d*_*LRR*_ = 14.5, *p* < .001; A: *R* = 0.846, *d*_*LRR*_ = 98.3, *p* < .001; Figure 7) and bimodal conditions (DS: *R* = 0.383, *d*_*LRR*_ = 12, *p* < .001; DA: *R* = 0.468, *d*_*LRR*_ = 18.7, *p* < .001; Figure 7), suggesting a substantial stability of sensory noise parameters across sessions. In other words, the majority of the variability in performance can be more readily explained by differences between individuals, rather than changes within individuals or measurement noise. Higher repeatability estimates in the auditory and disparity cues, compared to the size cue and the combined cues (DS, DA), can largely be attributed to the relatively higher variability between participants. That is, on average, size discrimination noise and combined cue noise was more similar between participants, while participants varied more strongly in the ability to discriminate depth based on audio or disparity. This can also be seen in the small sensory noise boxplots in Figure 7.

**Figure 7:**
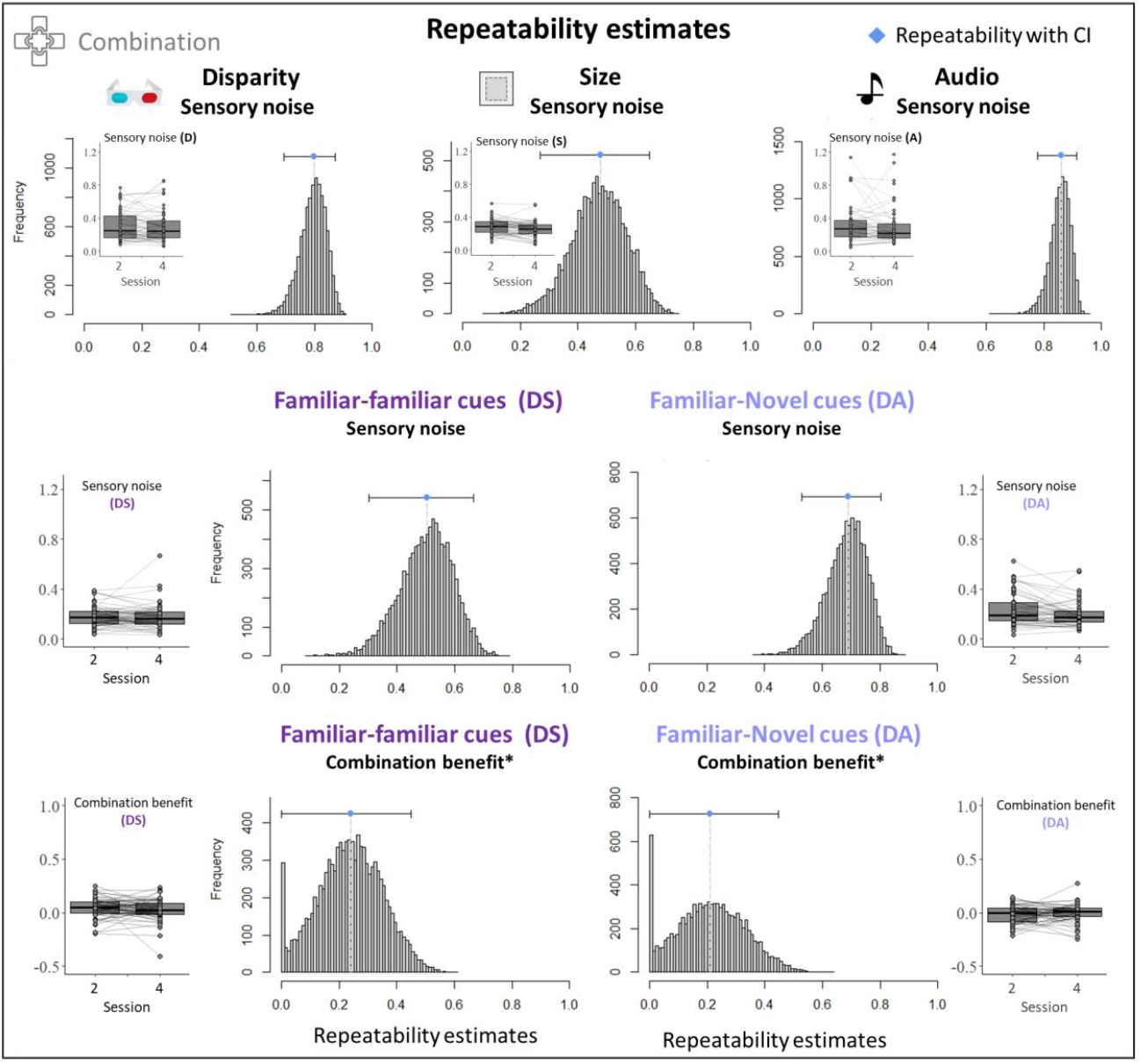
Within-individual repeatability was estimated for combination benefits (top row) and sensory noise values in the different bimodal (middle row) and single cue conditions (bottom row) across two sessions. Repeatability estimates are indicated as blue diamonds, error bars indicate CIs, based on 1000 or *10000 bootstrapped samples. Note that repeatability values of 0 in combination benefits were inflated by mixed model convergence failures. Frequency distribution shows sizable repeatability on converging models. Box plots that are presented within or next to repeatability estimate panels show the sensory noise and combination benefits for all participants in sessions 2 and 4, between which repeatability is estimated. Higher repeatability is reflected in a smaller within-participant variability, relative to between-participant variability. Sensory noise values are shown for each session and are connected within individuals.

Fitting combination benefits between the cue pairs led to several convergence failures in the mixed effect models, leading to an inflation of low repeatability value estimation. We therefore increased the number of bootstrap samples to 10000 and added permutation tests to serve as comparator for repeatability estimates. Nevertheless, repeatability measures of combination benefits should be interpreted with caution, given the inflation of low repeatability due to convergence issues. Combination benefits still showed a sizeable amount of repeatability within individuals across days (familiar-familiar: *R* = 0.239, *d*_*LRR*_ = 11.6, *p* < .001, permutation test: *p* = .029; familiar-novel: *R* = 0.213, *d*_*LRR*_ = 166, *p* < .001, permutation test: *p* = .056; Figure 7). Furthermore, there was no difference in the magnitude of combination between the sessions at the group level (familiar-familiar: *V* = 1298, *p* = .22; familiar-novel: *V* = 680, *p* = .338).

Taken together, the repeatability analyses suggest that not only sensory precision but also combination benefits can be reliably determined and are more stable within observers across time, than between observers. Hence, variability in sensory precision and perceptual precision benefits can be largely attributed to true inter-individual differences.

#### 3.3.2. Between-task correlations

To assess whether the degree of re-weighting and sensitivity to incongruence would predict combination performance, we cross-correlated these functional indices within each cue pairing. Overall, there was little evidence for intra-individual correlations between combination and re-weighting indices (*r* < 0.07; *p* > .66), nor between combination and incongruence sensitivity indices (*r* < 0.1; *p* > .50), suggesting that combination of the familiar-familiar and familiar-novel cue could not be explained by the degree of re-weighting nor by the sensitivity to congruence. As the re-weighting index depends on the initial weight applied to each cue, is based on *relative* weights (bound between 0 and 1) and corresponds to the change in weight based on the addition of a fixed amount of noise, this lack of correlation is not too surprising. Furthermore, the absence of a correlation between cue combination benefit in precision and sensitivity to congruence may suggest that participants may have used different decision strategies in the two tasks. Indeed, when a large conflict between the cues is noticed, observers might switch to following only one of the cues.

## 4. Discussion

Learning to integrate novel cues into our normal, perceptual repertoire underpins the flexibility of human cognitive and behavioural functioning in changing environments. Here, cue combination, an operation that takes place at the intersection of perception and cognition, provides a suitable candidate process via which novel sensory information can be efficiently integrated with already existing senses.

The present study assessed to what extent human adults can learn to integrate novel depth cues into their normal, perceptual repertoire with only minimal training. The processing of already existing, familiar cues was used as a baseline against which perceptual functioning with novel cues was compared. Perceptual functioning was assessed via markers that are typically observed with familiar cues: precision increases resulting from cue combination, dynamic re-weighting in line with sensory reliabilities, and sensitivity to cue mapping congruence.

Group-level analyses indicated that even after minimal training (∼300 trials), the novel auditory cue was already reweighted in line with sensory reliability and was responded to in a congruence-sensitive manner, similar to existing, familiar cues. However, in contrast to familiar cues, the novel audio cue was not combined with existing cues to depth (disparity) to produce a perceptual precision benefit. Instead, precision was, on average, more similar to the best, single cue. How can the absence of a precision benefit and a similar degree of re-weighting be reconciled? Crucially, while re-weighting may also result from alternative decision strategies, such as switching between the cues on an individual trial-level basis, a precision benefit cannot be explained by switching, but indeed, by true combination. As such, this suggests that, when presented with novel and familiar cues together, observers may have responded to noise changes by increasing the rate at which they based their judgements on the best, individual cue - rather than responding to noise changes by modifying a weighting with which they averaged (combined) the cues. The two familiar cues showed the precision improvement predicted by combination in line with statistical optimality. This contrasts with previous findings showing that cues to horizontal localization and simulated depth in VR can be quickly learnt and combined, albeit sub-optimally (Aston, Beierholm, et al., 2022; Negen et al., 2018).

Interestingly, while group-level results suggested that the novel cue was not combined with the familiar disparity cue, the present study uncovered strong inter-individual differences: about half of the participants showed a combination benefit with the novel cue; that is, they gained precision from having the familiar and novel cue available at the same time, compared to using only the best single cue. This benefit, as well as individual sensory noise estimates, were repeatable across different sessions, showing that a substantial part of the variability within sensory noise and combination benefit measures was attributable to variation between participants, rather than within participants. This suggests that the variability within the group was not a mere result of measurement noise or intra-individual variability over time. Instead, it suggests that some participants consistently combined the novel and familiar cues after only minimal training, while others did not. This may also offer an explanation as to why the group-level results of the current study contrasted those of previous studies what included between 10 and 12 participants in each group (Aston, Beierholm, et al., 2022; Negen et al., 2018). The present study comprised one of the largest samples in cue combination studies to date, and the multiple testing sessions allowed to probe its intra-individual stability. Notably, the stability of cue combination benefits would be line with an individually-specific tendency to bind the two cues (Körding et al., 2007; Shams & Beierholm, 2022). In other words, the belief that the two sensory cues arise from the same environmental feature (here: depth) may differ between individuals and mediate the combination of two cues. A stable causal inference belief, which has recently been shown by Odegaard and Shams (2016), may account for the variability between, and stability of cue combination benefits within individuals.

Contrasting how the two cue pairs (familiar-familiar, familiar-novel) were processed showed that re-weighting was not only expressed to a similar degree in the two cue pairs, but also that the number of participants that re-weighted cues in accordance with sensory noise changes was not dependent on the cue pair, with the majority of people re-weighting both cues, and only a small, equal number of people combining either of the individual cue pairs or none. What’s more, the observers’ empirically measured cue weights were significantly predicted by the relative sensory noise of the individual cues. This suggests that the sensory uncertainty of a novel cue can be instantaneously employed as cue weight and exemplifies the perceptual flexibility that this perceptual process offers for novel cue learning. This instant use of sensory precision in weighting is in line with previous findings in novel cue learning (Negen et al., 2018).

Even after minimal training, observers were sensitive to the congruence of the cue mapping that they were presented with. That is, when the cues were presented in their natural (familiar-familiar) or learnt (familiar-novel) cue mapping, they were discriminated with higher precision, compared to when they were presented with a mapping that was incongruent to the natural/learnt relationship. Here, the difference between congruent and incongruent precision was used as a marker of incongruence sensitivity and showed that the vast majority of observers were sensitive to the cue mapping, independently of the cue pair. Among the remaining participants, a larger number of participants were sensitive to the congruence of familiar-familiar cue pairs, compared to those of familiar-novel cue pairs. This suggests that the frequency with which a specific cue mapping is experienced may consolidate the automatic fusion of the two cues (Ernst, 2007). Alternatively, it may also suggest that noticeable incongruence between the cues leads to the adoption of different perceptual decision strategies, such as cue switching (Nardini et al., 2008; Scarfe, 2022).

While the majority of observers used the sensory uncertainty of the novel cue to weight sensory information and responded favourably to the congruence of the learnt cue pair mapping after minimal training, only half of them benefitted from combining the novel and familiar cues. In contrast, the two familiar cues were combined in an optimal fashion. As both processes were measured within the same individuals, it is unlikely that selection bias or inter-individual differences, which we showed to be present, were responsible for this lack of combination at group level. Indeed, more participants were likely to combine the familiar cue pair than the novel-familiar cue pair. One may wonder how the absence of a familiar-novel combination effect at group-level can be conceived – especially in light of previous studies finding (sub-optimal) benefits (Aston, Beierholm, et al., 2022; Negen et al., 2018). Firstly, it is possible that the minimal training that participants received (1-hour associative learning with feedback) was not sufficient for some individuals to consolidate causal inference beliefs that favour the integration of the novel cue mapping. In fact, an increase of sensory noise towards incongruently paired cues and optimal sensory re-weighting can also be explained by cue switching. However, neither cue switching nor following the best single cue could account for the precision benefits obtained by about half of the group’s observers. Hence, to better understand the effects that training duration and inter-individual differences have on the combination with novel cues, future studies need to employ longer and more comprehensive training paradigms and have the power to draw inferences at the individual level. Also, more large-sample studies are needed to better understand the factors that lead to the ability to combine novel cues instantaneously. Secondly, cue combination can be conceived as a process that consists of multiple hierarchical processing stages (Cao et al., 2019; Rohe et al., 2019; Rohe & Noppeney, 2015). As such, employing a multi-method approach to investigate the neural, behavioural, and phenomenological effects of novel cue learning (Nardini et al., 2024) can provide a more holistic understanding of the different stages of processing involved in the integration of novel sensory cues into the perceptual repertoire.

In conclusion, the present study shows that there is considerable variability between individuals in the uptake of a novel sensory cue: while a novel cue was immediately deployed for perceptual depth judgements, being weighted in accordance with its sensory reliability and mapping congruence, not all participants fully integrated it into native perception. Cue combination, a marker of natural perception that is evidenced by precision benefits in the combined cue estimates, was only evident in around half of the participants. While this may, at the group level, appear to be variability inherent to measurement imprecision, follow-up repeatability analyses suggested that this variability is likely a reflection of true inter-individual differences. In other words, our data do not suggest that participants fail to combine novel-familiar cues in general, but rather than some participants combine the cues, while others do not. Further in-depth psychophysical experiments that power the tasks to draw inferences at the individual level will be needed to corroborate these findings. Furthermore, the findings open up new questions about the factors that lead to inter-individual differences in the ability to integrate novel sensory cues into native perception. These findings will have implications for the training and uptake of sensory substitution and augmentation technology, as well as perceptual learning in general. Lastly, the present findings are the first to directly compare the integrative processing of familiar cues and newly learned cues. They thereby provide not only a direct benchmark of native perceptual functioning in depth perception, but also a platform for further investigations into the neural, perceptual, and subjective correlates of novel cue learning.

## Acknowledgements

This project has received funding from the European Research Council (ERC) under the European Union’s Horizon 2020 research and innovation programme (grant agreement No. 820185) and a Leverhulme Trust Research Project Grant (RPG 2017-097).

