## SupplementaryMaterials for "Learning new perceptual skills: Individual differences in the computations that integrate novel sensory cues into depth perception"

### 1. Deviations from pre-registration

#### 1.1. Index calculation:

In the pre-registration, we suggested calculating combination, re-weighting and incongruence sensitivity indices for each cue combination (XY) by quantifying the absolute effects in sensory noise reduction/enhancement and PSE shifts, relative to optimally predictable effects:

*Combination*:

$$CI_{XY} = \frac{\min(\sigma_X, \sigma_Y) - \sigma_{XY}}{\min(\sigma_X, \sigma_Y) - \sigma_{XY}^{opt}}$$

with $\sigma_X$ and $\sigma_Y$ being the sensory noise of the two individual cues, i.e. disparity and size or disparity and pitch; and $s_{XY}$ being the sensory noise of the combined cues. $\sigma_{XY}^{opt}$ indicates the sensory noise that would be expected based on optimal predictions (see equation 12).

*Re-weighting*:

$$RI_{DY} = \frac{(w_D^{emp} - w_{Du}^{emp})}{(d_{opt})} \quad \text{with } d_{opt} = w_D^{opt} - w_{Du}^{opt}$$

with $w_D^{emp}$ and $w_{Du}^{emp}$ being the relative weight placed on the disparity cue under non-noisy and noisy conditions, respectively. According to the optimal observer model, the predicted weights $w_D^{opt}$ and $w_{Du}^{opt}$ were based on the single sensory conditions and were estimated as the sensory noise of the disparity cue relative to the sensory noise of the other paired cue (see equation 8), under non-noisy and noisy conditions, respectively.

*Incongruence sensitivity*:

$$IS_{XY} = \frac{JND_Z - JND_{XY_{inc}}}{JND_Z - JND_{XY_{inc}}^{pred}}$$

Here, $JND_Z$ refers to the just noticeable difference (JND; equation 6) of the cue that participants followed, which could have been either of the two single cues *X* or *Y*, and was determined for each participant and cue pairing separately. $\mathrm{JND}_{XY_{inc}}^{pred}$ indicates the predicted JND for the incongruent condition for cues *X* and *Y*, and was derived from the weighted JND of the corresponding congruent condition $\mathrm{JND}_{con}$ via:

$$JND^{pred}_{XY_{inc}} = \left|\frac{1}{\left(2 \cdot w^{opt}_{X} - 1\right)}\right| \cdot JND_{XY_{con}}$$

The relative weight $w_X^{opt}$ for cue $X$ was derived from the single cue conditions for cue X and Y (see equation 8).

<u>*All indices:*</u>

As such, the combination index, re-weighting index, and incongruence sensitivity index provide measures of function relative to what would be expected following the optimal observer model. However, this measure is highly sensitive to the relative size of the possible maximal benefit and measurement noise (see Scheller & Nardini, 2023). That is, all empirically measured sensory noise values and weights are affected by a certain degree of measurement noise, inherent to the task as well as parameter estimation. Despite rigorous attempts to reduce measurement noise in the task, all parameter estimates were subject to measurement noise. As possible benefits (i.e., differences between the best single cue and the predicted optimal cue) are small, the effects of measurement noise in the single cue conditions (which are used to estimate the optimal prediction) are amplified.

For instance, while true indices of combination should theoretically vary between 0 (participant follows the best single cue) and 1 (participant integrates cues optimally), only around half of the sample showed integration indices between ±2. As expected, the noise ratio between the two individual cues strongly predicted the absolute magnitude of the combination index (familiar-familiar: $\beta$ = 5.25; $r$ = 0.59, $p$ < .001; familiar-novel: $\beta$ = 51.13; $r$ = 0.844, $p$ < .001), suggesting that the combination index may be more sensitive to the size of the maximally possible benefit than the actual degree of combination.

As a result, we decided not to report the indices proposed above, as scaling them by a noisy measure may inflate the contribution of the less relevant sensory cue noise ratio. Instead, we reported in the main text:

*CI*: the difference between the best single cue and the combined cue noise (equation 9)

*RI*: the change in disparity weight in response to added noise (equation 10)

*IS*: the change in sensory noise as a result of sensory incongruence (equation 11)

1.2. <u>Cue comparator for incongruence sensitivity assessment (fusion)</u>:

In the preregistration, we introduced our marker of 'incongruence sensitivity' as 'fusion'. This was based on the assumption that decreases in sensory precision (increases in noise) were reflective of forced fusion. However, we noted post data collection that the 2AFC task that we employed did not allow to arbitrate between fusion, which assumes the cues to be integrated automatically, and cue switching, whereby participants rely on individual cues at a time. Instead, the measured marker is more indicative of the sensitivity with which participants respond to the learnt stimulus mapping congruence.

However, to test for incongruence sensitivity, we need to employ a different comparison cue. Rather than comparing sensory noise of the incongruent trials to those of the best individual cue, it was compared to that of the congruent cue pairing. That is, while providing the same amount of sensory cues, with the only difference being cue congruence, we can directly measure how sensitive participants are to the congruence of the familiar or learnt cue mapping.

To assess congruence sensitivity of cue pair X and Y, we asked whether $\sigma_{XY}^{inc} > \sigma_{XY}$, rather than $\sigma_{XY}^{inc} > \sigma_{Z,XY}$, where Z is the cue that participants followed mostly during the judgements.

#### 1.3. Subjective Experience Measures

We pre-registered assessing changes in subjective experience, both quantitatively in terms of the realness/strength with which a percept of depth is elicited with the different cues, the level of effort that participants required to process depth with the used cues, and qualitatively, by asking participants about the strategy they employed. These data form the basis of a separate line of studies that extend our understanding of the subjective experiences of depth judgements within our tasks.

### 2. Power estimation

Power-estimation was based on simulations to quantify the probability of detecting a significant cue combination benefit (decrease in sensory noise of the bimodal cues relative to the best unimodal cue) under different possible parameter constraints, determined in a series of pilot studies. These parameters are the average sensory noise of the best cue, as well as the sensory noise ratio of both cues. As both parameters constrain the maximal benefit that an optimal observer could gain from combining the cues, and hence, possible effect sizes, they need to be taken into consideration when estimating power (Scheller & Nardini, 2023).

The power analysis outlines the probability for detecting a true cue combination effect at the group level, when all participants exhibit a true combination effect (Figure S1). One thousand simulations were run for each set of parameter constraints that varied around the estimated averages derived from pilot data from 6 participants, including: a minimum sensory noise of the best cue of 0.1, 0.2, 0.3 and 0.4, a ratio of sensory noise values of 1, 1.5, 2, and 3, and maximum individual lapse rates of 10%. Each set of parameter constraints was simulated for different participant group sizes. Data from 6 pilot participants indicated average sensory noise of the best cue to be 0.212 for familiar-familiar, and 0.164 for familiar-unfamiliar pairings, while the ratio of the best to worst cue was, on average, 1.55 for familiar-familiar, and 2.18 for familiar-novel cue pairings.

The power analysis showed that an average best sensory cue noise of 0.2, and a sensory noise ratio of 1.5 (based on pilot data for familiar-familiar combination), we would achieve 80.2% power with a sample of 30 participants. At the same time, a sensory noise ratio of 2 (based on pilot data for familiar-novel combination) offered 65.5% power with 60 participants. Hence, in order to reduce the cue noise ratio and to increase the chances of detecting a true combination effect, the sound cue increment was adjusted to match the average sensory noise of the visual disparity cue.

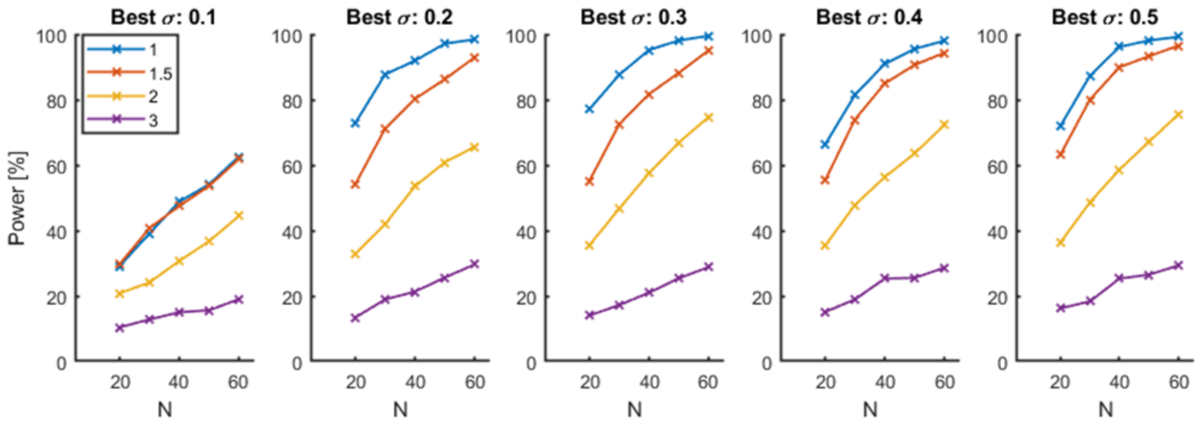


**Figure S1:** Power, defined as the probability of finding a true cue combination effect, as a function of sample size, the best sensory noise level, and sensory noise ratios between the cues.
